# Seagrass arabinogalactan-proteins: Are they important for adaptation to the marine environment?

**DOI:** 10.1101/2025.01.03.631252

**Authors:** Lukas Pfeifer, Kim L. Johnson, Antony Bacic, Thorsten B. H. Reusch, Carlos M. Duarte, Elizabeth A. Sinclair, Birgit Classen

## Abstract

During the late cretaceous period several lineages of angiosperm plants transitioned from land to the sea by successfully adapting to life in salt water, forming the polyphyletic group of seagrasses. Today, four seagrass families inhabit coastal systems and are deeply intertwined with health and welfare of these ecosystems. Adaptation to the ocean environment included changes in the composition of plant cell walls and associated glycoproteins. We have asked the question whether or not there is a convergent and similar arabinogalactan-protein glycan repertoire in all seagrasses, given initial findings of arabinogalactan-proteins with unique features in the well-studied eelgrass, *Zostera marina*. We isolated and characterized arabinogalactan-proteins from seven species covering the four major seagrass families using carbohydrate analysis and glycan immunoassays, along with a bioinformatic search for relevant gene pathways in newly published seagrass genomes and transcriptomes. Glycan parts of all seagrass arabinogalactan-proteins shared a high proportion of 1,4-linked glucuronic acids and terminal 4-*O*-methyl glucuronic acid residues. Trait-based dendrograms generated to inform phylogenetic-relatedness showed there was no phylogenetic signal among seagrass families and arabinogalactan-protein features. Transcriptomic datasets from *Cymodocea nodosa* and *Thalassia hemprichii* growing under hypersaline conditions showed an upregulation of enzymes involved in 4*O*-methylation and glucuronic acid transfer. We therefore conclude that environmental factors, especially salinity with higher monovalent ion concentration, influence seagrass arabinogalactan-proteins structure more intensely than phylogenetic history.

## Introduction

The polyphyletic group of seagrasses – marine angiosperm plants – evolved in Tethys seas of the Late Cretaceous 100-66 MYA (million years ago, Larkum et al., 2018) and successfully adapted to life in ocean waters. Seagrasses consist of four families (Cymodoceaceae, Hydrocharitaceae, Posidoniaceae and Zosteraceae). All seagrass families appear to share either a common land plant or freshwater ancestor of alismatid monocots (Ma et al., 2024), although adaptation to the marine environment occurred via three to four independent colonization events (Les et al., 1997; Waycott et al., 2018).

Seagrasses are distributed across shallow coastal environments around all continents, except Antarctica, with the highest species richness in tropical Indo-Pacific and south-west Australia (Short et al., 2007). Some families are only present in specific regions, such as the monotypic family Posidoniaceae (genus *Posidonia*) limited to the Mediterranean Sea and Australian coast regions (Aires et al., 2011). Seagrasses form extensive single or multispecies meadows, which are some of the most productive ecosystems on Earth, acting as globally significant carbon sinks (Duarte et al., 2005; Duarte et al., 2013). They also provide important habitats and/or feeding grounds for other organisms, including threatened animals like green turtles, manatees and dugongs (Unsworth et al., 2022). Despite their ecological importance and need to restore lost and damaged seagrass meadows, the knowledge on how seagrasses physiologically adapt to the saline environment remains limited (Sandoval-Gil et al., 2023; Olsen et al., 2016). A role for seagrass cell walls in the adaptation process is most likely. Yet, cell wall composition of seagrasses is still not well explored (Pfeifer and Classen, 2020a; Hoegh-Guldberg et al., 2014; Röthig et al., 2023).

Similar to cell walls of terrestrial angiosperms, cellulose and hemicelluloses are the main structural components in seagrasses (Davies et al., 2007; Syed et al., 2016; Ismael et al., 2023). However, seagrass cell walls present additional unique features, such as the presence of sulfated galactans in some species (Aquino et al., 2005; Kolsi et al., 2016) and unusual pectic macromolecules called apiogalacturonans (Gloaguen et al., 2010; Lv et al., 2015; Pfeifer, 2023). These consist mainly of galacturonic acid (GalA*p*) and apiose (Api) and have recently been shown to be a common feature of seagrasses worldwide (Pfeifer et al., 2022b). While lignin pathways are remaining in some species, their abundance is much reduced (Pfeifer et al., 2022b; Kaal et al., 2018; Ismael et al., 2023). Hypolignification is thought to enhance the flexibility of aboveground structures allowing stems and leaves to move with currents and waves (Ma et al., 2024).

Plant cell walls contain (glyco)proteins, especially arabinogalactan-proteins (AGPs), which belong to the group of hydroxyproline-rich glycoproteins (HRGPs, Silva et al., 2020). AGPs consist of a protein backbone and carbohydrate attached by an O-glycosidic linkage between hydroxyproline (Hyp) and a corresponding galactosyl-residue (Ellis et al., 2010; Seifert, 2020; Strasser et al., 2021). The carbohydrate part can be 90 to 99% (Nothnagel, 1997; Ma and Johnson, 2023) of the total mass and is dominated by the monosaccharides arabinose (Ara) and galactose (Gal). In land plants, galactan consists of a core chain with 1,3-β-D-Gal*p*, sometimes branched at O-6 to 1,6-β-D-Gal*p* side-chains, being then further decorated by terminal-, 1,3-α-L-Ara*f*, as well as 1,5-α-L-Ara*f* residues (Seifert and Roberts, 2007; Seifert, 2020; Strasser et al., 2021). AGPs have been implicated in cell signaling and have a variety of molecular functions including pollen development and sexual reproduction (Pereira et al., 2015; Pereira et al., 2016; Coimbra et al., 2007) and calcium-binding associated oscillatory processes (Lamport et al., 2014; Lopez-Hernandez et al., 2020). Recently, the presence of AGPs in seagrasses was demonstrated for *Zostera marina* (Pfeifer et al., 2020). An unusual structural feature of this AGP was its highly charged nature caused by 1,4-D-glucuronic acid (GlcA*p*) and terminal 4-O-methyl-GlcA*p* (4-O-MeGlcA*p*) residues, which may play a role in salt adaptation.

AGPs are widespread among seagrass species and this study aimed to elucidate the presence of AGP protein backbones and glycosyltransferase biosynthetic genes in published seagrass genomes and transcriptomes. We collected representatives across the four seagrass lineages (Cymodoceaceae, Hydrocharitaceae, Posidoniaceae and Zosteraceae) to investigate whether all species contained AGPs. Seagrass AGPs were isolated and characterized using biochemical and immunocytochemical methodologies. Bioinformatic searches for AGP protein backbones and glycosyltransferase biosynthetic genes in seagrass genomes broadened our knowledge on seagrass AGPs and helped to understand whether AGP structures might be related to seagrass phylogeny. Finally, gene expression data of *Cymodocea nodosa* and *Thalassia hemprichii* under salinity stress (Malandrakis et al., 2017;

Shen et al., 2022) were evaluated with regard to possible changes in biosynthesis or loss of AGPs. Altogether, these biochemical and bioinformatic data shed light on the potential role of seagrass AGPs in adaptation to the marine environment.

## Results

### Yields of seagrass arabinogalactan-proteins (AGPs)

AGPs were extractable from the aqueous extracts of all plant samples investigated and confirmed the presence of AGPs across seagrass species. Extracts of high-molecular weight polysaccharides and glycoproteins (including AGPs) from seven seagrass species gave yields between 1.4 % to 2.5 % relative to dry weight (Table S1), with the highest yields obtained for *Enhalus acoroides* and *Nanozostera noltii* (2.5 % in each species). AGP yields of the different seagrass species ranged from 0.02 to 0.10 % (w/w dry weight, Table S1).

### Protein moiety of seagrass AGPs

The AGP protein moieties of all investigated seagrass species show overall similarity to other angiosperm AGPs but also show characteristics which differentiate them from their non-seawater relatives. Hydroxyproline (Hyp) is the secondary amino acid responsible for the *O*-glycosidic linkage between protein and carbohydrate moieties in AGPs. Hyp content can therefore be used to evaluate the state of glycosylation in an AGP sample was determined spectrophotometrically (Figure 1A and B). All investigated AGPs had Hyp values between 0.59 to 0.93 % (w/w; Figure 1 left). Notably, both *Posidonia* species show the highest amounts of Hyp, while all other AGPs were more similar with Hyp values between 0.59 and 0.69 % (w/w).

**Figure 1.**
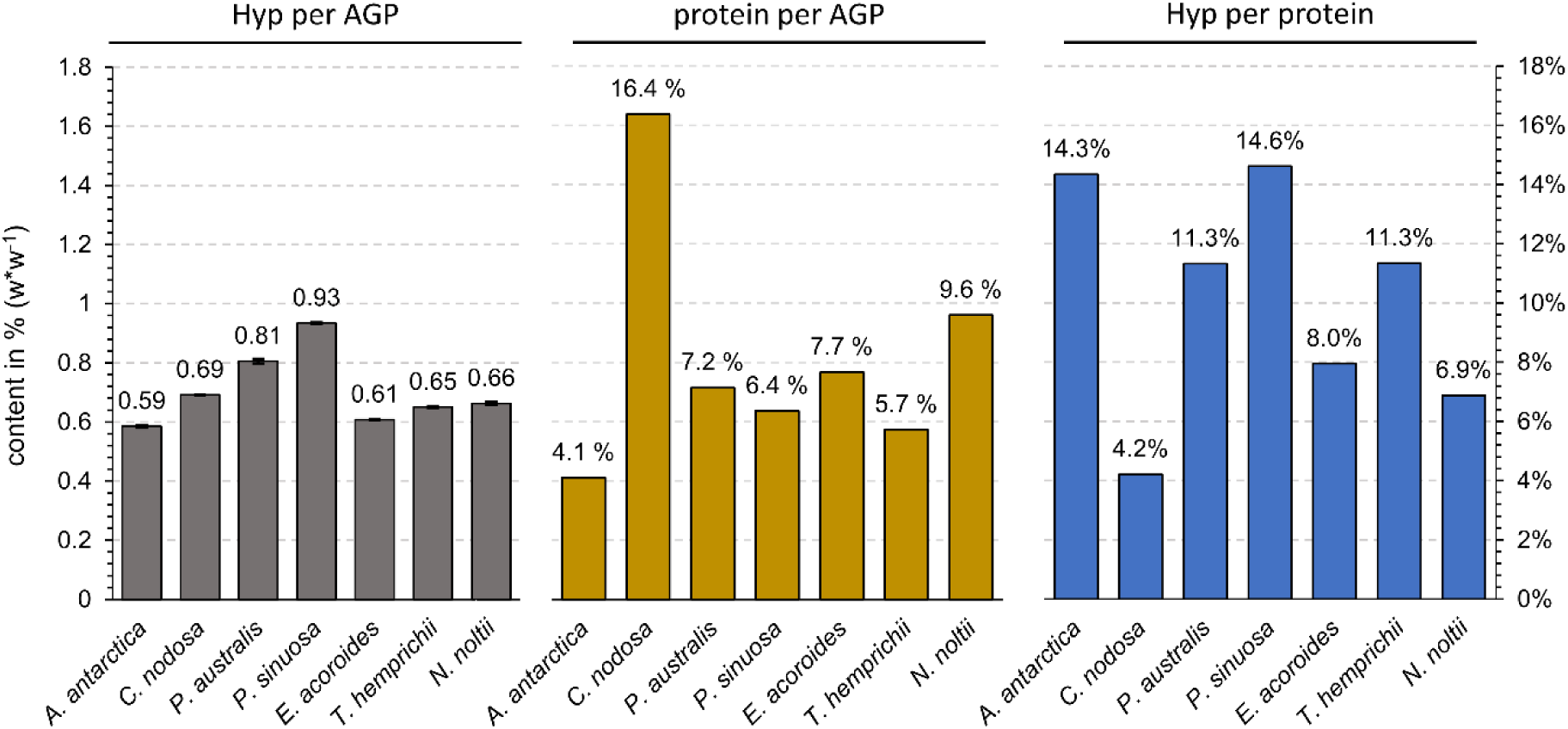
Determination of hydroxyproline (Hyp) and protein contents. All values are given in % (w/w^-1^). Hyp content (left) was quantified colorimetrically (n = 3, error bars represent standard deviation), protein content (middle) was inferred from elemental analysis (N x 6.25) (n = 1). Hyp per protein (right) was calculated based on the two other determinations.

The total protein contents of AGPs varied between 4.1 and 16.4 % (Figure 1, middle) – a value that is (except for *C. nodosa*) in the range usually found in angiosperms (1-10 %; Fincher et al., 1983; Nothnagel, 1997). Figure 1 (right), shows the amount of Hyp relative to the protein content, which ranged between 4.2 and 14.6 %.

AGP protein sequences were identified in all seagrass genomes (available for *Thalassia testudinum*, *Amphibolis antarctica*, *Cymodocea nodosa*, *Posidonia australis*, *Posidonia oceanica*, *Zostera marina* and *Zostera muelleri*) and transcriptomes of the four families (Figure 2A). The genome of the closely related alismatid freshwater plant *Potamogeton acutifolia* was included for comparison.

**Figure 2.**
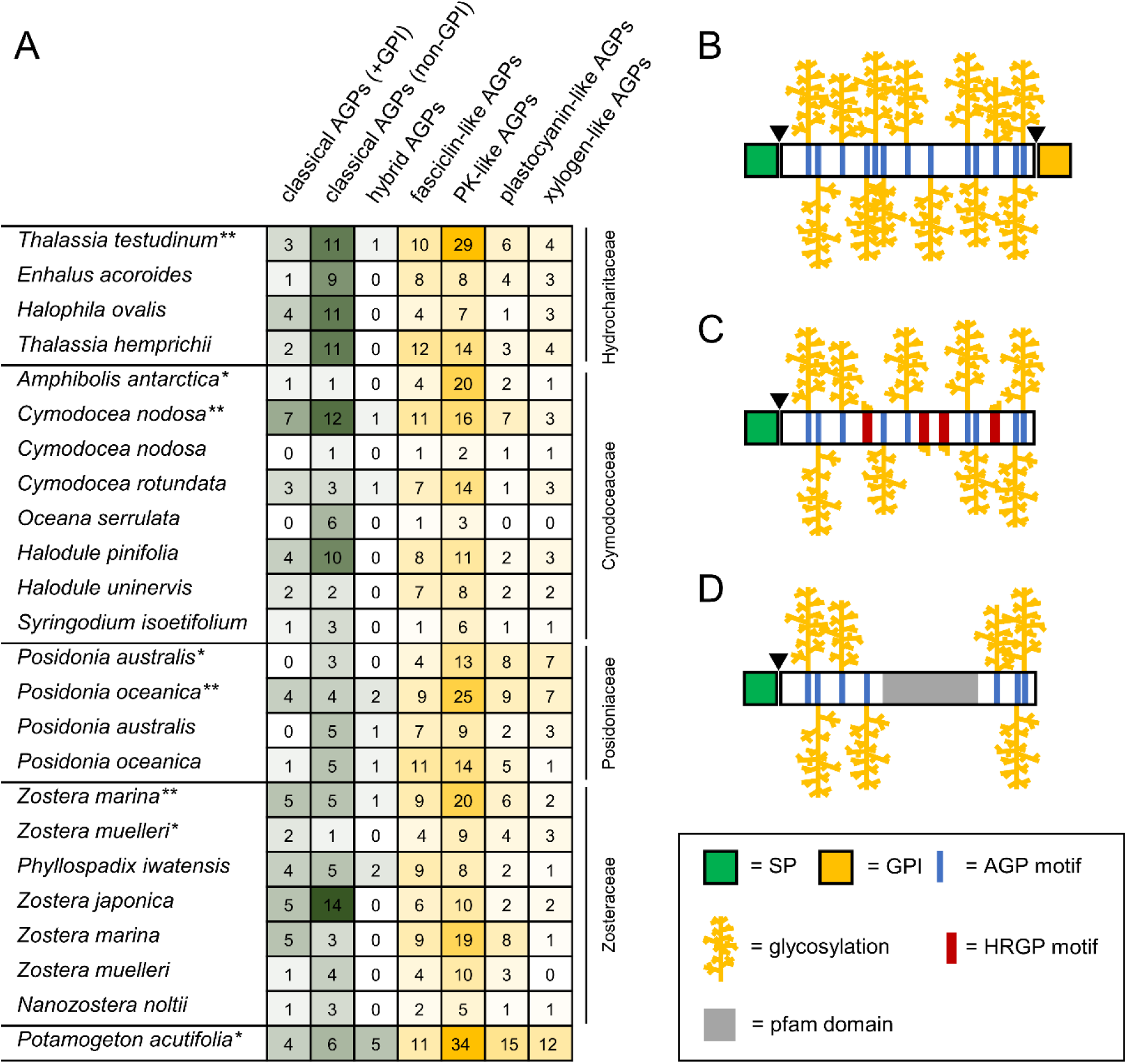
Bioinformatic search for AGP backbone sequences in different seagrass genomes (*: draft genomes; **: chromosome-level genomes) or transcriptomes. A) Number of sequences in the AGP classes in different species of the four seagrass families; B, C, D) Schematic representations of classical (B), hybrid (C) and chimeric (D) AGPs.

These AGP sequences can be classified into classical AGPs (Figure 2B) with typical amino acid motifs rich in “PAST” (proline, alanine, serine, threonine), the hybrid AGPs with a mixture of AGP and other HRGP motifs (Figure 2C) and chimeric AGPs containing additional functional protein domains (Pfam domains; Figure 2D).

Sequences for classical AGPs (with and without GPI-anchor, i.e. class 1 and 4) were identified in all seagrass samples (Figure 2A). In most samples, numbers for classical AGPs with GPI-anchor (class 1) were lower compared to those without GPI-anchor (class 4).

The total number of classical AGPs in seagrasses was comparable to *P. acutifolia*, although the proportion of classical AGPs without GPI-anchor was higher in the Hydrocharitaceae, *C. nodosa*, *Halodule pinifolia* and *Zostera japonica* (Figure 2A). Class 5-8 comprises hybrid AGPs with AGP bias and other motifs of HRGPs, e.g. class 5 has additional motifs typical for cross-linked extensins and class 6 lacking cross-linking motifs (Johnson et al., 2017a). The number of hybrid AGP sequences identified was either low (1 - 2) or completely absent in seagrasses, whereas five sequences were identified in *P. acutifolia*. For chimeric AGPs, we focused on the most common Pfam domains (fasciclin-like, protein kinase-like, plastocyanin-like and xylogen-like) known to occur in chimeric AGPs from angiosperms. The number of chimeric AGPs in seagrasses was lower for all types relative to *P. acutifolia*, especially the plastocyanin-like and the xylogen-like AGPs. Higher numbers of plastocyanin-like and xylogen-like AGPs were detected in the Posidoniaceae. For some species, genome and transcriptome sequences were available (*C. nodosa, P. australis*, *P. oceanica*, *Z. marina*, *Z. muelleri*). These are sometimes quite similar, e.g. in *Posidonia* species and *Z. marina*, but in case of *C. nodosa*, the number of AGP sequences in the transcriptome was very low in comparison to the genome.

### Carbohydrate moiety of seagrass AGPs

Beside monosaccharides characteristic for all AGPs, seagrass AGPs show higher amounts of GlcA in 1,4-linkage and as terminal 4-*O*-methylated residues. The compositional analysis revealed dominance of Gal and Ara in AGPs of all species with molar percentages ranging from 40.1 - 47.0 % and 27.7 - 44.7 %, respectively (Figure 3A and B, Table S2). The composition of the AGP from *E. acoroides* was distinguished with the highest content of Ara, the lowest level of Rha and an absence of Fuc. It also had the highest Ara: Gal ratio at 1: 0.9 in comparison to the other species (between 1: 1.3 and 1: 1.6). GlcA residues were present in all AGPs primarily as GlcA, but also with low levels of 4-O-Me-GlcA (Figure 3A). The presence of 4-O-MeGlcA in all samples seems to be characteristic for seagrass AGPs. Other monosaccharides of seagrass AGPs were Rha in amounts between 3.5 and 11.8 % (Table S2), with exception of *E. acoroides*, which contained only traces of this monosaccharide, as well as Glc, Man, Xyl and Fuc in small amounts.

**Figure 3.**
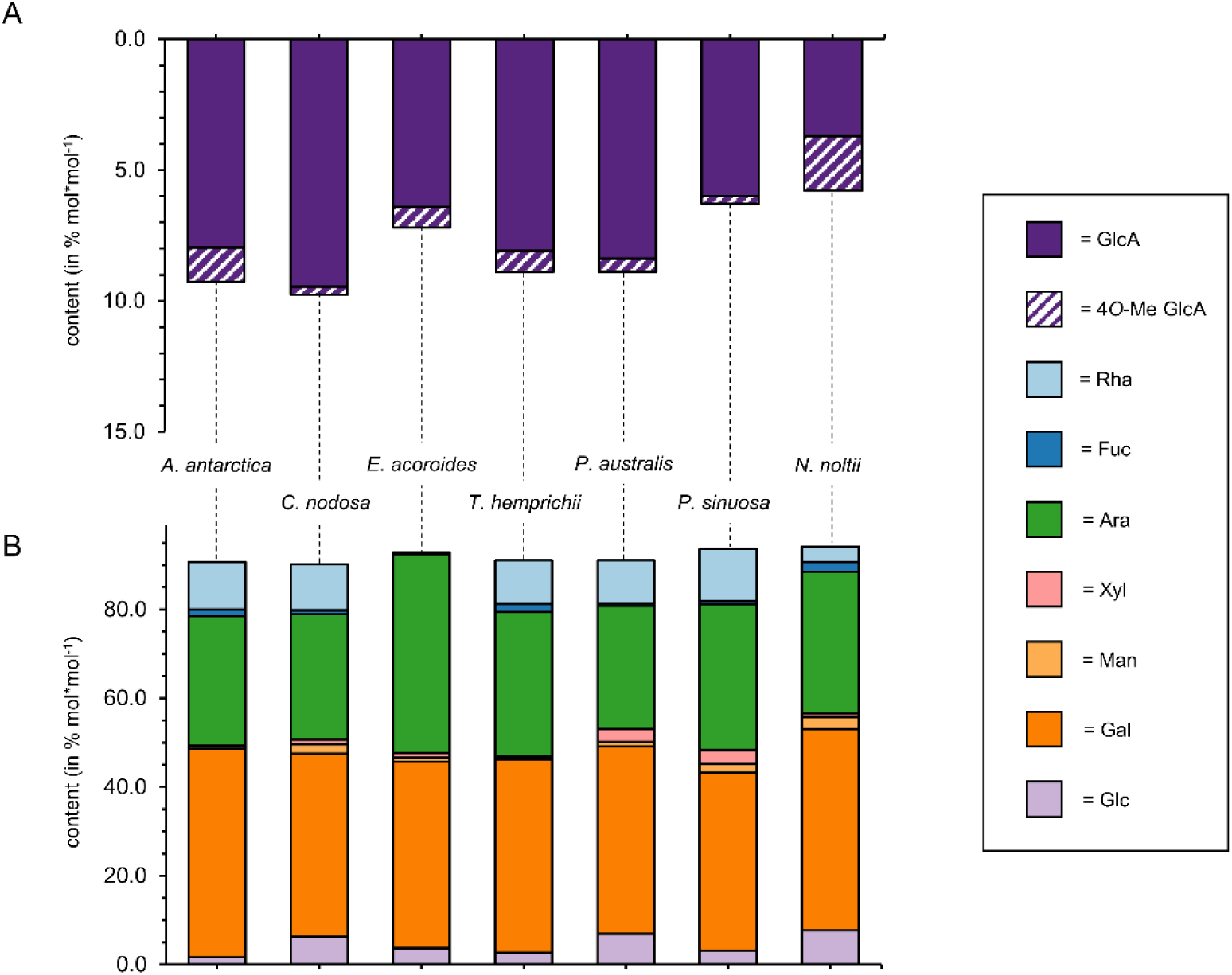
Monosaccharide composition of seagrass AGPs. A) Content of uronic acids (% mol mol^-1^) determined according to (Kim and Carpita, 1992). B) Neutral monosaccharide composition determined by gas-chromatography (% mol mol^-1^). See also Table S2.

Uronic acid reduced samples were submitted to methylation analysis for linkage determination (Table 1). The galactan core reveals the typical structure of a type-II-AG with β-D-Gal*p* in 1,3-, 1,6- and 1,3,6-linkage. The strong dominance of the 1,3,6-linkage type in all species (26.9 – 34.5 %) is striking and indicates highly branched carbohydrate. The side chain structures were heavily decorated with additional sugars, as oxalic-acid treatment increased the relative amount of 1,6-Galp in comparison to the non-treated sample (Figure S1, Figure S2). There were low numbers of small and unbranched side chains as indicated by low 1,6-Gal*p* contents prior to acid degradation (Figure S2B). This number increased after degradation with oxalic acid. The proportion of 1,3-Gal*p* remained stable after acid degradation thus supporting a branched side-chain structure with many modifying monosaccharides

**Table 1.**
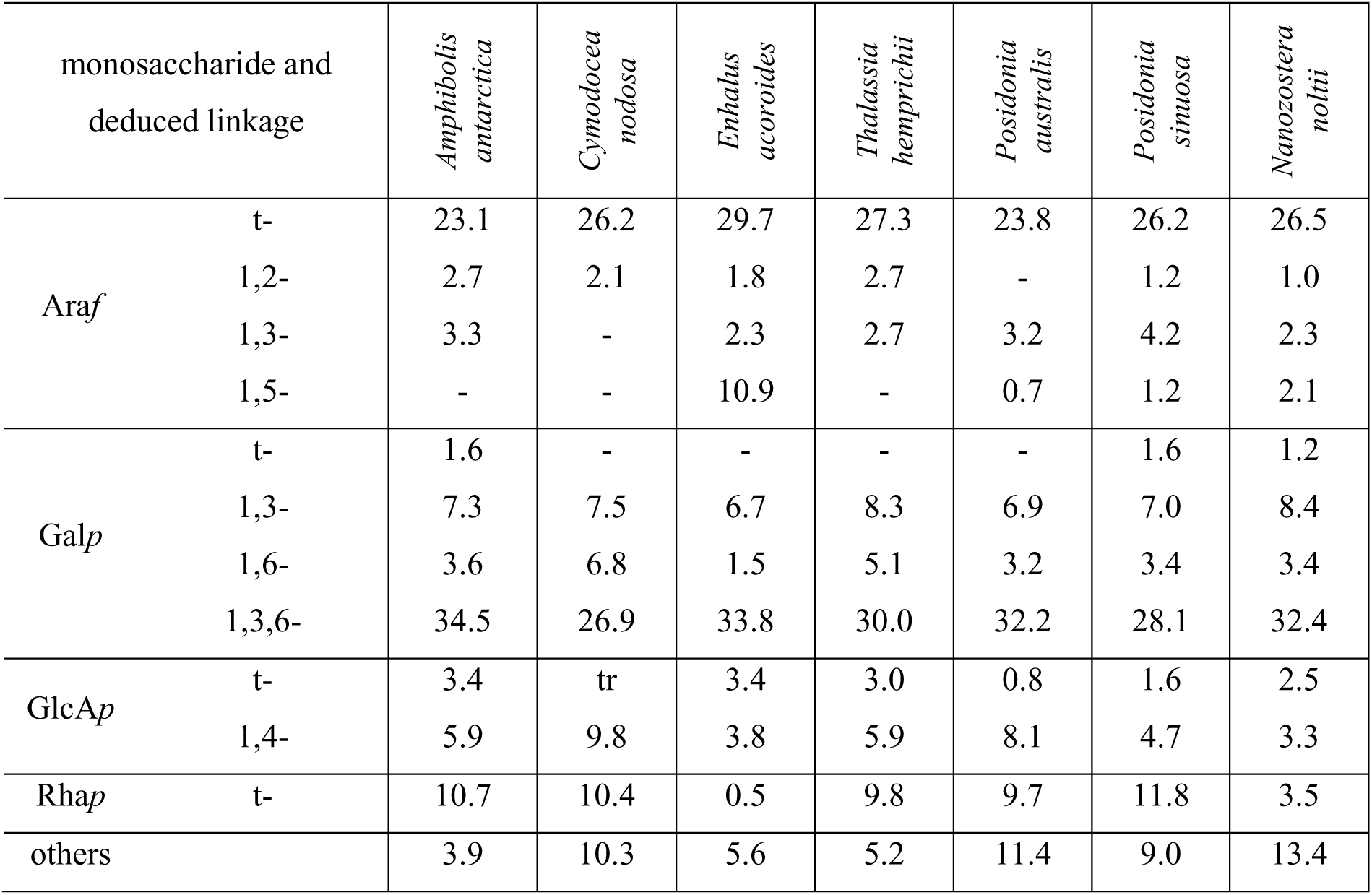
Monosaccharide linkage-type analysis (% mol/mol) of seagrass AGPs.

(Figure S2C). Ara*f* is the main decorating monosaccharide (see also Figure 3) and present mainly as a terminal residue. 1,5-Ara*f* was present in four species, but only *E. acoroides* was characterized by a high incidence of this Ara linkage. All seagrass AGPs were rich in GlcA, present as terminal and 1,4-GlcA residues. It is important to note that the *t*-GlcA value will also include the terminal 4-*O-*MeGlcA as these cannot be differentiated in a standard methylation analysis. For further distinction of both residues, methylation with deuterium-labeled iodomethane (as performed by Pfeifer et al., 2023) would be necessary but the proportions can be deduced directly from the monosaccharide analyses (see Table S2). The ratio of terminal to 1,4-linked GlcA was nearly 1:1 in *E. acoroides* and *N. noltii* and shifted more to the non-terminal residue in all other seagrasses, particularly in *C. nodosa* and*P. australis*. Longer chains of GlcA were implied with a ratio tending to 1,4-linked residues. Rha*p* was only present as a terminal residue.

### Detection of AGP-carbohydrate epitopes by antibodies

Monoclonal antibodies that detect specific AGP glycan epitopes combined with enzyme-linked immunosorbent assays (ELISA) were used to determine the fine-structure of the isolated seagrass AGPs (Figure 4). JIM13 and LM2 reacted to all seagrass species in their highest concentrations. The most intense reaction for JIM13 was detected with *C. nodosa* AGP, followed by *N. noltii* AGP, *P. australis* and *P. sinuosa* AGP, and moderate interaction in the remaining seagrass AGPs. JIM13 is a classical AGP-glycan antibody, but the epitope structure is not fully understood. A trisaccharide consisting of β-GlcA-(1,3)-α-GalA-(1,2)-Rha has been shown to bind to this antibody (Knox et al., 1991; Yates et al., 1996), but neither 1,4-GlcA nor 1,2-Rha were detected in the linkage analyses of seagrass AGPs (see Table 1). Therefore, other carbohydrate motifs must be responsible for binding, probably including Rha and GlcA.

**Figure 4.**
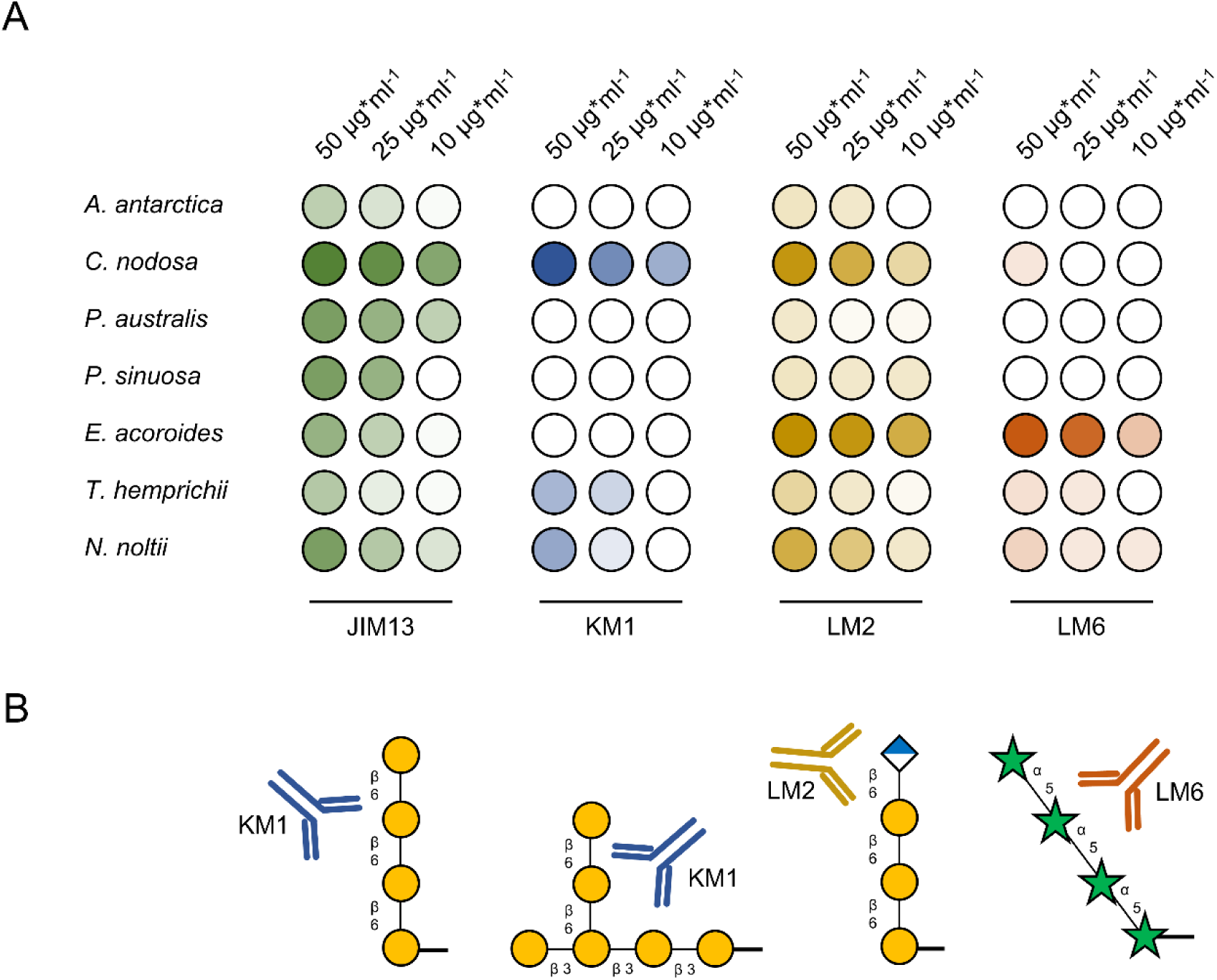
Results of the ELISA experiments with antibodies directed against AGP glycan-motifs. A) Heatmaps for seven seagrass species with the four selected antibodies JIM13, KM1, LM2 and LM6. B) Described epitopes of the antibodies represented according to Ruprecht et al. (2017) and Willats et al. (1998). Monosaccharides are symbolized by yellow circles (Gal), green stars (Ara) and blue-white squares (GlcA).

*C. nodosa* reacted most intensely to the monoclonal antibody KM1, while *T. hemprichii* and *N. noltii* showed low intensities and all others lacked reactivity. LM2 and LM6 bound strongly to *E. acoroides* AGP and moderately to AGPs from *C. nodosa*, *T. hemprichii* and *N. noltii*. LM2 is an AGP-antibody, which detects terminal GlcA attached to 1,6-Gal side chains (Figure 4B) and was originally generated against rice AGPs (Smallwood et al., 1996; Ruprecht et al., 2017). LM6 was raised against pectic polysaccharides and binds to 1,5-α-L-arabinans (Figure 4B; Willats et al., 1998). AGP from *E. acoroides* was rich in 1,5-linked Ara (Table 1) and consistent with the presence of this glycan motif.

### Classification of AGP-related carbohydrate-active enzymes (CAZys)

We searched for proline-4 hydroxylases (P4Hs), for glycosyltransferases (GTs) and for glycosylhydrolases (GHs) to shed light on biosynthesis and reconstruction of seagrass AGPs. P4Hs and GTs are responsible for biosynthesis of AGPs, namely hydroxylation of proline and subsequent glycosylation, while GHs are involved in possible remodelling of AGP carbohydrate fine structures. Figure 5A depicts our proposed structure of AGP in *Zostera marina* (Pfeifer et al., 2020) in combination with results of our bioinformatic search for the enzymes (Figure 5B; Dataset S1-S4). The described enzyme activities of GTs and GHs are highlighted (Figure 5A). P4Hs were present in all genomes and transcriptomes revealing that the first step providing reactive hydroxyl groups for *O*-glycosylation is possible in seagrasses. The responsible GTs belong to different GT families that have been classified in the carbohydrate active enzymes (CAZy) database (http://www.cazy.org/). Known members of GT families, involved in AGP biosynthesis, were found in all seagrass species. GALT29A homologs were missing in some species, while all relevant GT31 clades were represented in the dataset. All seagrass sequences contained some members of the GT14 family of glucuronosyltransferases (GlcATs), except the *Oceana serrulata* transcriptome. Most abundant were homologs to GlcAT14A+B. Homologs were detected in all seagrass sequences, except in the *A. antarctica* and *P. australis* genomes. GT37 family members were detected in all seagrass transcriptomes and genomes with an overall higher presence in the Cymodoceaceae. The β-arabinofuranosyltransferase RAY1 was present in low numbers or missing in the investigated seagrass genomes and transcriptomes. Beside GTs, some known GHs, such as AGAL1-3 (GH27), RsBGAL1 (GH35), RsAraf (GH3) and members of GH43 and GH79 are important for AGP metabolism. Sequences of these degrading enzymes were present in all seagrass families. Homologs of RsAra*f* were especially abundant in the Hydrocharitaceae (genera *Thalassia* and *Enhalus*).

**Figure 5.**
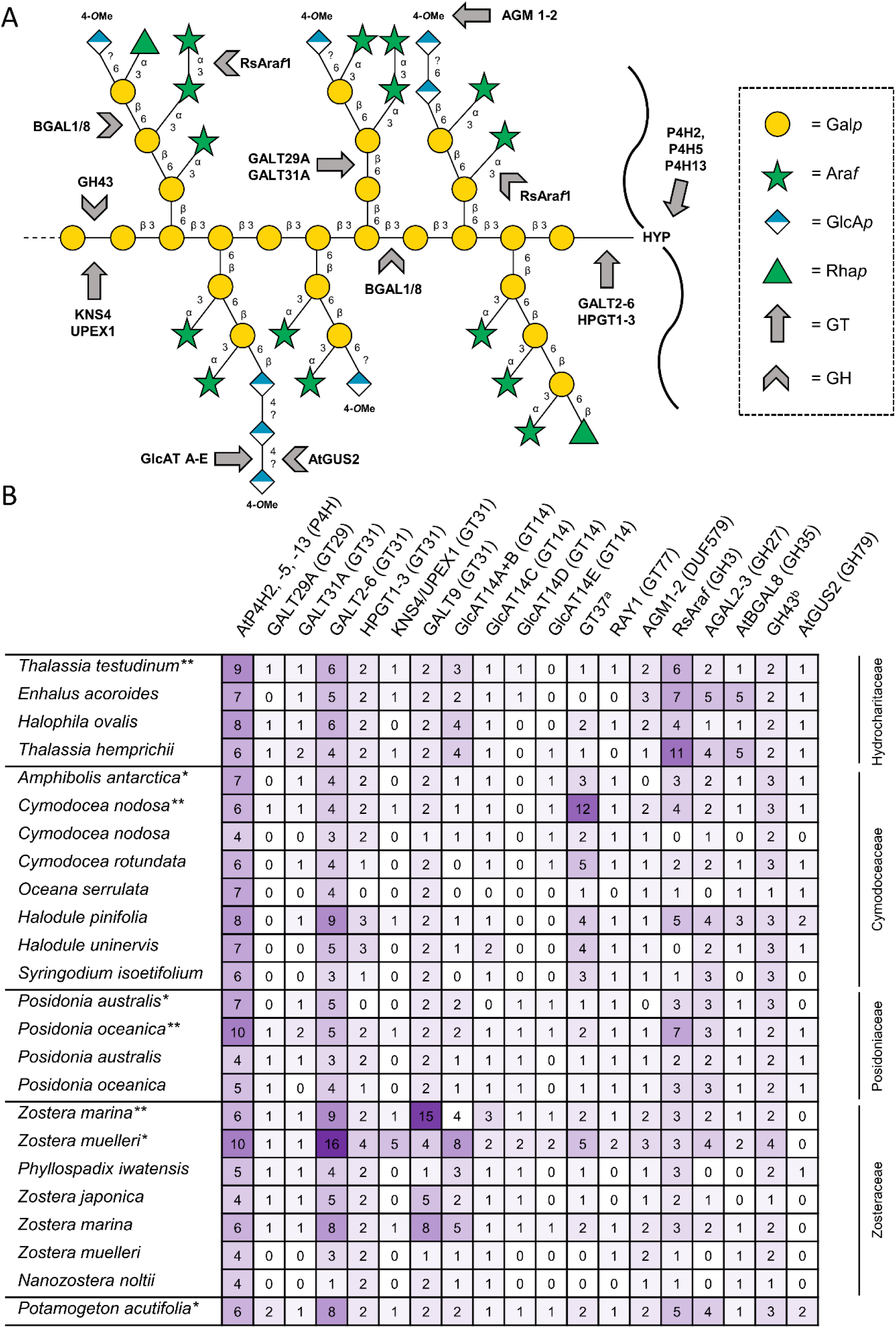
AGP-related carbohydrate-active enzymes (CAZys) in different seagrass genomes (*: draft genomes **: chromosome-level genomes) or transcriptomes. A) Structural proposal for AGP from *Zostera marina* (Pfeifer et al., 2020) showing enzymes predicted to hydroxylate proline and synthesize carbohydrate; note that for *Zostera marina* AGP all terminal glucuronic acids were found 4-*O*-methylated, this is not the case for all investigated species. B) Numbers of predicted CAZys sequences by species from four seagrass families (see also Dataset S1-S4).

### Differentially expressed genes in relation to salt stress

Transcriptome data for two seagrass species grown under hyperosmotic salt stress revealed some cell wall-related CAZys likely taking part in stress reaction (Figure 6). There was no overlap in downregulated CAZys between *C. nodosa* and *T. hemprichii*. In contrast, enzymes belonging to families GT8 and DUF579 were upregulated during salt stress in both species.

**Figure 6.**
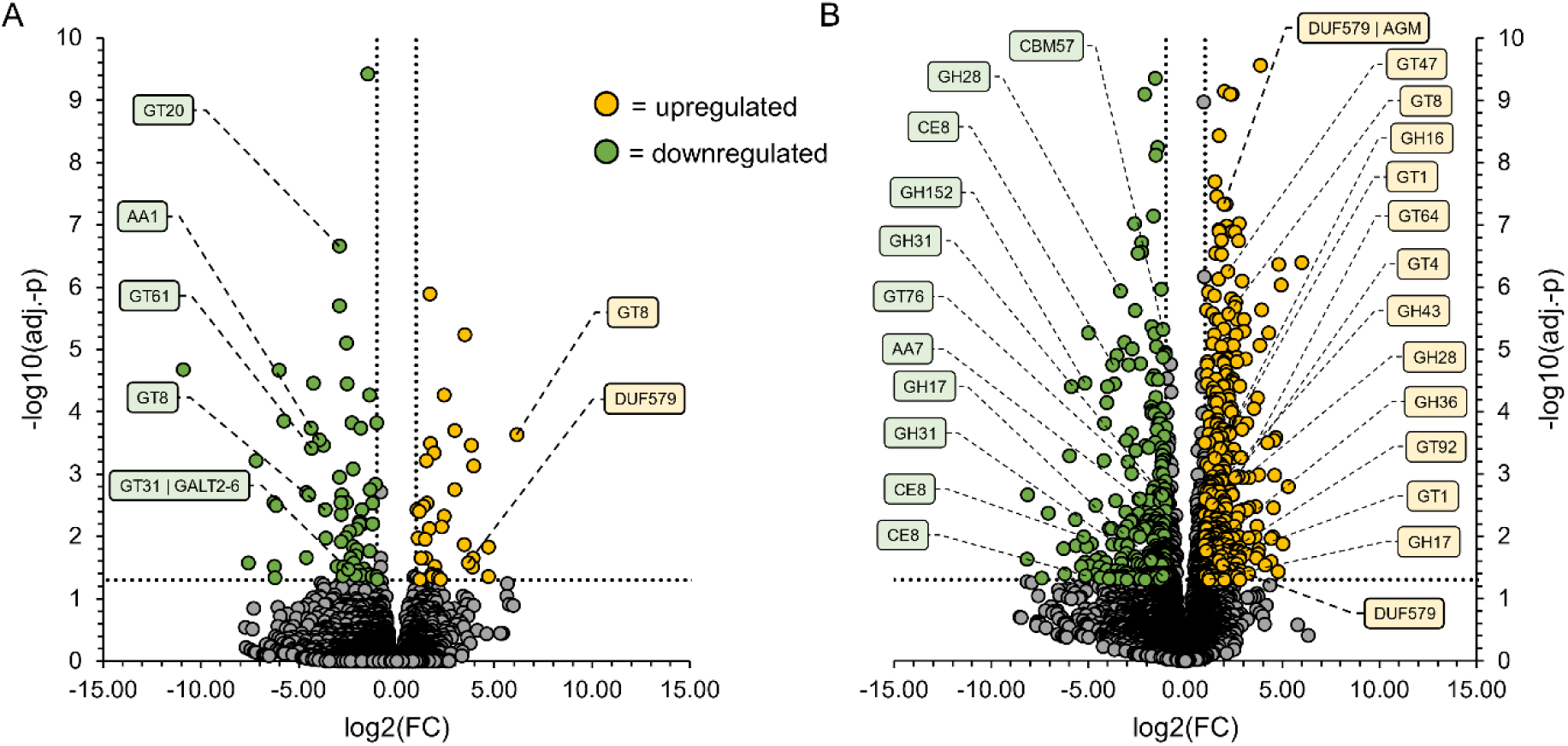
Volcano plots of the differentially expressed genes in response to elevated salinity. All predicted CAZymes are annotated. The colors represent up- or downregulated genes for A) *Cymodocea nodosa* and B) *Thalassia hemprichii.* Dotted lines indicate significance at 2- and 0.5-fold change (FC), as well as an adjusted *p*-value of 0.05.

CAZy genes were also upregulated under salt stress in *T. hemprichii* and comprised members of the families GT47, GT1, GT4 and GT64, as well as glycosylhydrolases of the families GH16, GH17, GH28, GH36 and GH43. Most of the GT47 members in plants identified, to date, are involved in biosynthesis of different cell wall polysaccharides including pectic GlcA transferases (Iwai et al., 2002; Zhong and Ye, 2003) and 1,3- and 1,5-α-AraTs. Upregulated genes of the families GT1 and GT64 were involved in different types of plant stress responses (Li et al., 2017; Lin et al., 2022). Family GH16 includes cell-wall-modifying transglycanases, e.g. xyloglucan endotransglucosylase (Franková and Fry, 2013). Members of GH43 have been characterized as Golgi-located exo-β1,3-galactosidases acting on cell wall-bound AGPs in *Arabidopsis* (Nibbering et al., 2020). GH28 comprises structurally related enzymes mainly known from fungi that hydrolyze pectins, including polygalacturonases, rhamnogalacturonases and xylogalacturonases (Sprockett et al., 2011), and GH36 include α-galactosidases acting on gum Arabic AGPs (Sasaki et al., 2021). Finally, members of GH17 have been identified in *Vitis vinifera* which are associated with response to abiotic stress including salt stress (Pervaiz et al., 2021).

## Discussion

Our overall findings reveal that AGPs are ubiquitously present in the cell walls of seagrasses across all phylogenetic lineages. We found conserved protein and core glycan properties while the fine-structural features diversified in the investigated seagrass species. Genes encoding enzymes involved in the biosynthesis of a common structural feature, namely 4*O*-MeGlcA, were found upregulated in hypersaline conditions when looking at transcriptomic data. All of this highlight evolution of an adaptive strategy in seagrass cell walls to cope with saline conditions.

### Seagrass AGPs show conserved protein and core glycan structures

Bioinformatic searches for AGP protein backbones in seagrass genomes and transcriptomes confirmed the presence of classical AGPs, with and without GPI anchors, in all seagrass families. The gene numbers were comparable to *Potamogeton acutifolia*, a freshwater monocot of the closely related alismatid family of Potamogetonaceae, which is in favor of a more or less conserved AGP protein repertoire in Alismatales. The number of protein backbones for classical AGPs in seagrasses was slightly lower than in other monocots and core eudicots, with an average of 5 GPI-anchored AGP and 8 non-GPI-anchored AGP sequences. For example, 17 GPI-anchored AGPs and 12 non-GPI-anchored AGPs were detected in the *Arabidopsis genome*, while 14 GPI-anchored AGPs and 20 non-GPI-anchored AGPs were present in the *Brachypodium* genome (Johnson et al., 2017a). There were higher numbers of non-GPI-anchored AGPs in seagrass families, a trend also observed in other monocot and dicot transcriptomes (Johnson et al., 2017b), as well as fern genomes (Mueller et al., 2023). The sometimes-occurring divergence between genomic and transcriptomic data, e.g. number of GPI-anchored AGP sequences in *Cymodocea nodosa*, highlights the need for more well-curated genomes to better understand molecular mechanisms in a specific plant group. Basing general conclusions about presence/absence of genes on the genomic information is more reliable. This is reflected when looking at the BUSCO scores (Figure S3) as a metric for genome/transcriptome quality. In case of *C. nodosa* the missing and fragmented genes in the BUSCO analysis are much higher. It has to be taken into account that most of the investigated transcriptomes were *de novo* assembled (Chen et al., 2022) before the publication of the well-curated genomes of Cymodoceaceae, Hydrocharitaceae and Posidoniaceae (Ma et al., 2024). A reference-guided *de novo* approach (e.g. as proposed by Lischer and Shimizu, 2017) would be an option to further disentangle the cases in which higher sequence numbers were detected in the transcriptomes. Nevertheless, the absence of most sequences in the transcriptomes could be a hint for only minor contribution of those under natural conditions.

All seagrass families were characterized by the occurrence of chimeric AGPs, i.e. proteins containing AGP motifs plus a functional protein domain. We focused on well described chimeric AGPs of seed plants like protein-kinase (PK)-like, plastocyanin-like, fasciclin-like (FLA) and xylogen-like AGPs (Basu et al., 2016; Costa et al., 2019; He et al., 2019; Johnson et al., 2003; Ma et al., 2017; Shafee et al., 2020), which were all present in seagrass genomes and transcriptomes. The PK-like AGPs were most abundant and comparable to our previous investigations on *Zostera marina* (Pfeifer et al., 2020) and ferns (Mueller et al., 2023), underlining their importance, although their functions are still unclear. These chimeric AGPs are proposed to work as sensor molecules in plants and brown algae (Hervé et al., 2016).

The carbohydrate moieties of seagrass AGPs reveal unique structural features, in contrast to the protein moieties, which are generally comparable to angiosperm land plants. Nevertheless, it has to be taken into account that there might be differences in architecture of the protein domains like e.g. described for fasciclin-like AGPs (Shafee et al., 2020). The galactan backbones of seagrass AGPs share typical characteristics of land plant AGPs with a 1,3-linked β-D-Gal*p* backbone highly branched at the 6-position (1,3,6-linked) to side chains of 1,6-linked β-D-Gal*p* (see core galactan structure features found in GC-MS of oxalic-acid treated samples, Figure S1 and S2). Members of GT29 and GT31, responsible for synthesis of the type-II-galactan framework (Silva et al., 2020) were present in all species, although homologs of GALT29A were missing in many cases. Homologs of GALT2-6 were most abundant. In *Arabidopsis*, these enzymes are responsible to transfer the first Gal onto Hyp, the first step of AGP galactosylation (Basu et al., 2015a; Basu et al., 2015b). RsBGAL1 (GH35) is a β-galactosidase from *Raphanus sativus* and hydrolyzes β-1,3- and β-1,6-Gal residues (Kotake et al., 2005). In *Arabidopsis* AtBGAL8 is the closest homolog to that enzyme. Presence of these homologs in all investigated seagrass genomes hint towards the maintenance of the galactan scaffold as it is known from land plants (Leszczuk et al., 2023). These findings underline a conservation of the galactan backbone in all four seagrass families, with modifications in the decorating sugars. It has been shown previously that chain length in the 1,6-linked β-D-Gal*p* side-chains contributes to important physiological processes, for example cell elongation and cellulose deposition (Kikuchi et al., 2022). While the proportion of the chain length was conserved in five investigated angiosperms (Ito et al., 2020), we see here that seagrass AGPs tended to have smaller but more side chains (as indicated by the amount of 1,3,6-linked Gal*p*). Kikuchi et al. (2022) connected long side chains with increasing cellulose deposition activity. The latter was an influential factor to plant cell wall elasticity. This might indicate that seagrass cell walls are stiffer due to their reduced and branched AGP side chains. Contribution of AGPs to cellulose deposition was shown in different studies (Tan et al., 2012; Nibbering et al., 2022; MacMillan et al., 2010). Beside this biomechanical influence, the glycan parts were functionally active mostly *via* their minor monosaccharides attached to the galactan side chains. The calcium binding through β-glucuronidation (Lopez-Hernandez et al., 2020) or the pollen tube guidance function of 4O-MeGlcA attached to 1,6-linked Gal*p* in *Torenia* (Mizukami et al., 2016) can be mentioned in this direction. This exemplifies that different physiological functions are influenced by various molecular aspects of AGP (glycans) (Kikuchi et al., 2022). Although they are generally present in minor amounts, the monosaccharides attached to the galactan side chains – the outer surface modification – should get attention in the light of molecular functionalities.

### AGP glycan surface modifications show unique characteristics in seagrasses specific to marine plants

AGP glycan surface modification in seagrasses shows biochemical differences to land plant AGPs as uronic acids are a dominant structural feature. Differences in AGP side-chain structure were apparent in *Enhalus* AGPs with higher amounts of Ara*f* compared to the other seagrass AGPs. Knowledge on arabinosyltransferases (AraT) is poor and β-AraT RAY1 is the only AraT described to be working on AGPs to-date (Gille et al., 2013). AGPs contain α-1,2-, 1,3- or 1,5-linked Ara*f* residues, so the role of RAY1 in AGP arabinosylation is questionable (Showalter and Basu, 2016).

The numbers of homologs of the α-L-arabinofuranosidase RsAra*f* (GH3, Kotake et al., 2006) in the Hydrocharitaceae (Figure 5) seem to be contradictory to higher amounts of Ara*f*, especially in *Enhalus* AGP. Nevertheless, the higher numbers of degrading enzyme homologs (GH3, GH27, GH43; Kotake et al., 2006; Imaizumi et al., 2017) suggest a general capability to modify Ara residues in response to environmental factors. An increase in homolog numbers of RsAra*f* could therefore suggest its importance to maintain (net balance against similarly high numbers of AraT) or modify (possessing the response capacities to degrade the Ara*f* rapidly following an environmental stimulus) the Ara contents. The cell wall composition of *Enhalus* featured higher apiosyl and arabinosyl abundance compared to all other species (Pfeifer et al., 2022b). Interestingly, *Enhalus acoroides* is the only seagrass species which does not complete its lifecycle fully submerged as pollination occurs at the water surface. The male flowers detach and float on the water until then reach a female flower where pollination can occur, pollen and styles remain dry (Ackerman, 2006).

Family GT37 contains fucosyltransferases (FucTs) and a high number of sequences belonging to this family were found in the genome of *Cymodocea nodosa*. It is likely that these enzymes are involved in biosynthesis of other polysaccharides, as our biochemical studies only detected traces of Fuc in seagrass AGPs.

Substantial amounts of β-D-GlcA*p* in terminal position and in 1,4-linkage were biochemically detected in all investigated seagrass species, in contrast to land plant AGPs that generally feature lower amounts of GlcA*p*. Our analysis was not able to identify the seagrass specific enzymatic basis for that, as no GT responsible for 1,4-linkages of GlcA in AGPs is described. Terminal GlcA residues are seen as an interplay between the biosynthesis and the remodeling process. On the biosynthesis side, the GT14 family includes different GlcATs which add GlcA to Gal chains of AGPs (Knoch et al., 2013; Dilokpimol and Geshi, 2014). Comparison of homolog numbers in the main clades (GlcAT14A-E, see Dataset S4) revealed family-specific dominances (high homolog numbers in GlcAT14A+B in Zosteraceae and Hydrocharitaceae), while the general GT14 repertoire was present in all seagrass species. De-glucuronosylation by β-glucuronidases (GH79 family member AtGUS2, Eudes et al., 2008) seem to be possible as most seagrass species contain homologs of the respective enzyme. Strikingly, the recently updated version of the *Z. marina* genome completely lacks AtGUS2 homologs (Figure 5). This was also observed in other Zosteraceae members, with the exception of *Phyllospadix iwatensis*. The combination of high numbers of GlcAT homologs and lack of AtGUS2 homologs in Zosteraceae might be a strategy to maintain a high level of GlcA in AGPs. Multiplication of the GlcAT homologs could therefore be a mechanism to guarantee this important functionality.

Methylation of GlcA in *Arabidopsis* AGPs is performed by AGM1+2 belonging to the DUF579 family (Temple et al., 2019). Most, but not all, seagrass genomes and transcriptomes contained similar AGM1+2 homologs (Figure 5). Given the fact that our biochemical data showed presence of 4*O*Me-GlcA in all seagrass AGPs in the range of 3.5 to 10.8 % (mol mol^-1^), it is likely that there are other methyltransferases (from the DUF579 family or in another yet unknown family) working on AGPs. Due to the current lack of genetic transformation strategies in seagrasses, comparable studies showing differential activities in organs of known *Arabidopsis* AGMs (Smith et al., 2020) are not currently possible.

### Features of seagrass AGP glycans did not correlate with phylogenetic position

There was a mismatch between established phylogenetic positions (Les et al., 1997; Waycott et al., 2018) and trait-based dendrograms (Figure 7). This mismatch supported the hypothesis that environmental factors were responsible for architectural variation in seagrass AGP glycan. This should not question the currently accepted phylogeny, as the seagrass positions are supported by multiple studies (Les et al., 1997; Waycott et al., 2018; Ma et al., 2024; Chen et al., 2022; Da Silva et al., 2021; Tuya et al., 2024). The two commonly used sequences *rbc*I and *mat*K are reliable to resolve phylogenies on seagrass genus and species level, respectively (Papenbrock, 2012). Tuya et al. (2024) showed strong phylogenetic signal caused by phylogenetic niche conservation for seagrasses. The authors summarized that early adaptation processes of the respective ancestors of the three independent seagrass lineages to a very similar environment during the late Oligocene and early Miocene influenced trait evolution much more that later species radiation. Genomic comparison of different seagrass lineages supported a habitat-driven convergent evolution characterized by common morphological and physiological features, e.g. lack of stomata, gas exchange through permeable cuticles, effective osmoregulation and presence of aerenchyma in roots and rhizomes (Wissler et al., 2011; Lee et al., 2018; Ma et al., 2024). In general, convergent evolution is prevalent among plant families that adapted independently to similar extreme environments (Xu et al., 2020).

**Figure 7.**
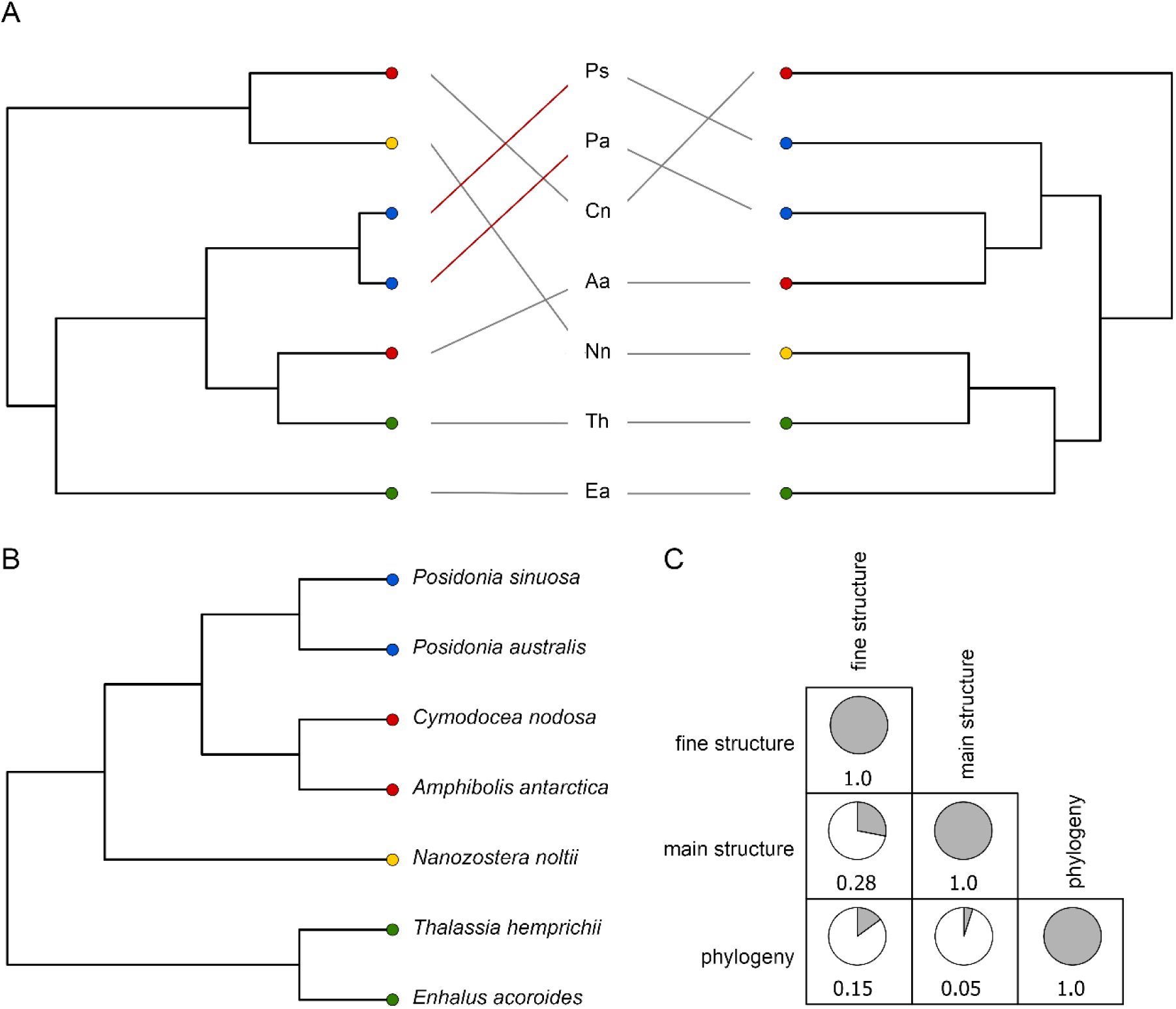
Similarity cluster dendrograms based on seagrass AGP analyses. A) Tanglegram representation based on ELISA results with 4 antibodies (fine structure, left side) and combined monosaccharide composition and linkage composition data (main structure, right side). B) Relationship of the four seagrass families based on *mat*K and *rbc*L, redrawn from Waycott et al. (2018). C) Cophenetic correlation matrix of the main and fine structure dendrograms with the current phylogeny. Phylogenies appear monophyletic in the absence of other aquatic families.

### Adaptation of seagrass cell walls to salinity

Marine brown and red algae are phylogenetically much older (Denoeud et al., 2024; Borg et al., 2023) and only very distantly related to green plants including marine angiosperms, however, a look at their cell walls might nevertheless be instructive for understanding the required biochemical features of cell walls under high sodium and chloride ion concentration. Marine algae possess cell walls which contain highly charged polysaccharides like the alginates and fucose-containing sulfated polysaccharides of brown algae (Mazéas et al., 2024) and the sulfated galactans of red algae (Ciancia et al., 2020). Sulfated galactans also occur in the marine angiosperm *Ruppia maritima* and their amount correlates with environmental salinity (Aquino et al., 2005). In the same study, it was shown that cultivation of rice in presence of NaCl also stimulates biosynthesis of carboxylated polysaccharides in cotton (Zhong and Läuchli, 1993) and rice (Aquino et al., 2011). Obviously, acidic polysaccharides (either sulfated or carboxylated) are needed to cope with external NaCl. Pectins represent the major part of charged cell wall polysaccharides in seagrasses (Pfeifer et al., 2022b). Possible candidates that might be involved in seagrass cell wall adaptation to salt are apiogalacturonans (Pfeifer, 2023; Pfeifer et al., 2022b), unusual pectic polysaccharides which crosslink via borate. Borate crosslinking is quite stable in the presence of sodium, so apiogalacturonans might be an advantage under salty conditions (Ma et al., 2024). A key enzyme in the biosynthesis of apiogalacturonans, UDP-D-Api/UDP-β-D-Gal*p*-xylose synthase, has expanded in seagrasses (Ma et al., 2024). A further candidate involved in handling salt stress is pectic homogalacturonan (HG), which is methylesterified in the Golgi and secreted into the wall in a highly methylesterified form. Pectin methylesterases catalyse the demethylesterification of HG in the cell wall, thus offering the possibility of fine-tuning electronegativity (Pelloux et al., 2007). Free carboxyl groups of HG are cross-linked via Ca^2+^ to form complexes called “eggboxes” (Grant et al., 1973), but under saline conditions, monovalent Na^+^ is able to displace Ca^2+^ and disrupt these structures, thus maintaining cell wall elasticity (Ma et al., 2024), an important feature to cope with currents and wave action.

Our biochemical and bioinformatic data point at an involvement of AGPs in the salinity-response strategy of seagrasses by high amounts of terminal GlcA residues which are partly modified by methyl ether groups on position C4. Lineage-specific variations in the side chain organisation and terminal sugar decoration of AGPs has also been observed in mosses and ferns, where the content of terminal Rha is higher and a special feature of bryophyte, fern and gymnosperm AGPs is *t*-3-O-MeRha (Fu et al., 2007; Bartels et al., 2017; Bartels and Classen, 2017; Baumann et al., 2021). This monosaccharide has not been detected in any AGPs or polysaccharides from flowering plants. Thus, the occurrence of methylated sugars in AGPs is not unusual in the light of cell wall evolution (Baumann et al., 2021; Mueller et al., 2023; Pfeifer et al., 2023; Pfeifer and Classen, 2020b; Pfeifer et al., 2022a). It might be speculated, that during land plant evolution, the nature of AGPs evolved from a more hydrophobic to a more hydrophilic surface, as Ara is more hydrophilic compared to Rha and methylation contributes to less hydrophilic AGP structures (Fu et al., 2007; Pfeifer et al., 2022a).

The evolutionary origins of AGPs are still enigmatic, as no AGP *sensu stricto* has been isolated from any algae. AGP protein backbone sequences are present in algae, and attempts to isolate AGPs resulted in molecules with atypical monosaccharide compositions (Eder et al., 2008; Přerovská et al., 2021). Recently, a rhamnogalactan-protein was isolated from cell walls of the freshwater alga *Spirogyra pratensis* (Zygnematophyceae) using Yariv precipitation that consisted of the galactan core structure typical for AGPs but decorated with Rha instead of Ara (Pfeifer et al., 2022a). For seagrasses, sharing a common ancestor with monocotyledonous land plants, we found that the AGP molecules are different with high amounts of GlcA residues resulting in a polyanionic surface. This is important with regard to adaptation to the marine environment. Comparable to GalA residues in pectins, pairs of two AGP GlcA molecules are ideal binding partners of Ca^2+^-ion crosslinking., The Ca^2+^ binding capacity in *Z. marina* AGPs was shown by microscopy and isothermal titration calorimetry and found to be in the range of calmodulin (Pfeifer et al., 2020). Other investigations revealed that uronic acid containing AGPs bind Ca^2+^ more strongly than does pectin (Lamport et al., 2020) and infer that Ca^2+^ is able to protect against damage caused by NaCl (Lahaye and Epstein, 1969; Cramer et al., 1985). Binding of Ca^2+^ to seagrass AGPs in the extracellular matrix possibly protects against high Na^+^-concentrations in the marine environment.

Transcriptome data from the seagrasses *C. nodosa* and *T. hemprichii* grown under salt stress support this hypothesis. Enzymes belonging to GT8 and DUF579 families were upregulated at elevated salinities hyper-osmotic in both species (Malandrakis et al., 2017; Shen et al., 2022). Members of the GT8 family are glucuronosyltransferases (GlcATs) responsible for the biosynthesis of terminal GlcA residues on glucuronoxylans (GXs, Rennie et al., 2012). Members of the DUF579 family are responsible for methylation of GlcA in GXs and AGPs (Temple et al., 2019). Both these studies (45 psu and 50 psu) exceeded the salinity preferences of most seagrass species (range 9-52 psu; Touchette, 2007), so this reaction may differ from physiological responses under naturally elevated variations in natural salinity levels, such as the metahaline in Shark Bay (Booth et al., 2022). Methylation of GlcA possibly leads to a subtle variation of the Ca^2+^ binding subunit which might support the discrimination of Na^+^ (Lamport and Várnai, 2013; Lamport et al., 2014). A microscopic study suggested a role of AGPs as carriers of Na^+^ from the apoplast to the vacuole during salt stress (Olmos et al., 2017), which helps to cope with osmotic pressure.

AGPs might be also involved in signaling, aside from the possible barrier function of GPI-anchored AGPs close to the plasma membrane Ca^2+^-ions released by AGPs in an auxin-dependent process might be responsible, leading to a lower periplasmic pH by regulation of the plasma membrane (PM) H^+^-ATPase (Lamport et al., 2014). Further possible mechanisms include involvement of oligosaccharides enzymatically cleaved from the AG glycan moieties or association of GPI-anchored AGPs with classical transmembrane proteins (Ma and Johnson, 2023).

Overall, seagrasses examined here showed a presence of AGPs across all phylogenetic lineages. They shared specialized glycan structural features with high amounts of uronic acids (+/-4-*O*-Me-GlcA) resulting in a highly charged surface. These electronegative surfaces are likely a convergent adaption of seagrasses to the marine environment. Pectins and AGPs with special features possibly act synergistically to cope with living in a marine environment. Both macromolecules were able to influence the balance of Ca^2+^ and Na^+^ in the extracellular matrix and are possibly involved in signaling cascades leading to further salt adaptation processes. Our data further strengthen the hypothesis that high levels of Na^+^ ions compete with Ca^2+^ ions in the marine environment thus making calcium-binding *via* charged-residues an important factor for maintaining calcium homeostasis (Lamport, 2024). The next step to enlighten this functionality would be the generation of knock-out mutants for a seagrass species. Unfortunately, aseptic laboratory culture and transformation of the now fully sequenced seagrass species (Ma et al., 2024) is currently a challenge. Shifting the focus of the mainly *Arabidopsis*-centered plant molecular biology field to a much more diverse set of model organisms (including seagrasses) from various plant (sub-)lineages would be a great stimulus for a more general understanding of the underlying mechanisms.

## Materials and methods

### Plant material

Seagrass samples from seven species were collected in the Red Sea (*Enhalus acoroides* and *Thalassia hemprichii*), Indian Ocean of Western Australia (*Amphibolis antarctica*, *Posidonia australis* and *Posidonia sinuosa*), Mediterranean Sea (*Cymodocea nodosa*) and Baltic Sea (*Nanozostera noltii*; Table S3). All plant material was cleaned from coarse pollution, rinsed in tap water and freeze-dried or oven-dried (below 60 °C).

### Isolation of high-molecular weight fraction (HMF) and purification of AGPs

A MF 10 basic grinder (IKA-Werke GmbH & Co.KG, Staufen, Germany) was used to prepare the dried seagrass samples for extraction. A sieve size of 1.0 mm was chosen and the powder was extracted according to Pfeifer et al. (2020). AGPs were precipitated using the specific dye β-glucosyl-Yariv (βGlcY) reagent (Yariv et al., 1967). The absolute yields of Ara and Gal, determined in the monosaccharide composition analysis were calculated and both values combined. The HMFs were dissolved in double-distilled water and an equal volume of βGlcY in sodium choride solution (0.3 mol l^-1^) was added. The mixture was incubated overnight at 4°C and a precipitate formed. The precipitate was then separated by centrifugation (19.000 *g*, 4°C, 30 min, Heraeus Multifuge X3R, Thermo Fisher Scientific Corp.) and redissolved in double-distilled water. The solution was heated to 50°C (IKA RET basic, IKA-Werke GmbH & Co. KG) and slowly sodium hydrosulfite/ dithionite (Sigma-Aldrich Corporation, St. Louis, MO, USA) was added until color change from red to white-yellow was achieved. This resulted in a final concentration of 1-5 % (m/v) hydrosulfite. After that, dialysis against water for 3 days at 4°C was performed (MWCO 12-14 kDa, Medicell Membranes Ltd., London, UK). The AGP fraction was then freeze-dried (Christ Alpha 1–4 LSC, Martin Christ GmbH, Osterode, Germany).

### Reduction of uronic acids and deuterium-labeling

For reduction of uronic acids in the AGPs, the samples were dissolved in double-distilled water and the procedure of Taylor and Conrad (1972) with slight modifications according to Pfeifer et al. (2020) was used.

### Acid hydrolysis of AGPs

Uronic acid reduced (UR) samples of the different seagrass AGPs were subjected to hydrolysis in 12.5 mM oxalic acid for 5 h at 100°C. The samples were dissolved in a concentration of 10 mg/mL in a Pyrotube C vial (Associates of Cape Cod Inc., East Falmouth, MA, USA) and placed in a heating block (Bioblock Scientific, Thermolyne Corp., USA). Afterwards, the hydrolyzed solution was precipitated in cold ethanol in a final concentration of 80 % (v/v).

### Determination of glycan composition

Monosaccharide composition analysis was performed using the procedure of Blakeney et al. (1983) with modifications in the hydrolysis step (2 M trifluoroacetic acid for 1 h at 120°C) and GC-analysis according to Pfeifer et al. (2020).

For determination of uronic acids, the quantification method of Kim and Carpita (1992) was used. Primary fragments of per-acetylated hexoses (m/z: 187, 217, 259, 289) and their 6,6-dideutero-labelled derivatives (+2 m/z) were detected in the uronic-acid reduced AGP samples by GC-MS measurements (GC parameters are stated below) in selective ion mode.

### Quantification of hydroxyproline and protein content

Hydroxproline (Hyp) content was determined photometrically following the procedure of Stegemann and Stalder (1967). A calibration with Hyp solutions in the concentrations 0.6 µg/mL, 1.2 µg/mL, 1.8 µg/mL, 2.4 µg/mL and 3.0 µg/mL was performed. A control with pure buffer instead of the oxidation reagent was performed for each sample to exclude the influence of other possible absorbing components. Elemental analysis was performed as described in Mueller et al. (2023).

### Linkage determination by *per*-methylation analysis

After uronic acid reduction the AGP, as well as additionally oxalic acid hydrolyzed AGPs were dissolved in dimethyl sulfoxide (DMSO). The dissolved samples were methylated according to Harris et al. (1984) by alternating addition of activated DMSO carbanion and iodomethane. Afterwards, the methylated samples were hydrolyzed, reduced and acetylated. The resulting partially methylated alditol acetates were extracted with dichloromethane and injected into GC (7890B, Agilent

Technologies, Santa Clara, USA; OV-1701 column, 25 m, 0.25 mm, 0.25 µm, Macherey-Nagel, Düren, Germany) with coupled flame ionization detection (FID) and mass spectrometry (MS) (5977B, Agilent Technologies, scan-range 40-500 m/z, electron ionization mode). The GC running parameters were used as described by Pfeifer et al. (2020). The internal standard *myo*-inositol was used.

### ELISA with AG-specific antibodies

Four different monoclonal antibodies (JIM13, KM1, LM2 and LM6, each in a dilution of 1:20) were tested in an ELISA for binding to the different seagrass AGPs. Three concentrations of the native AGPs (50 µg/mL, 25 µg/mL and 10 µg/mL in double-distilled water) were pipetted into the cavities of 96-well microplates (Nunc, Nalge Nunc International, Roskilde, Denmark) and dried at 37.5°C with open cover for 3 days. After that, the experiment was performed according to Pfeifer et al. (2020).

### Similarity clustering of the results

The results of the neutral monosaccharide composition, linkage type analysis, and ELISA results were imported into RStudio (version 1.2.5042) environment. After scaling and calculating an euclidean distance matrix, the results were hierarchical clustered (complete linkage method) and plotted as tanglegrams by the “dendextend” package (Galili, 2015). A cophenetics correlation matrix of these dendrograms was calculated and visualized by the package “corrplot” (Wei and Simko, 2024).

### Bioinformatic search for AGP backbones and AG-related enzymes

Eight seagrass genomes (Olsen et al., 2016; Ma et al., 2024; Lee et al., 2016; Bayer et al., 2022) and 16 seagrass transcriptomes (all from Chen et al. (2022) were searched for AGP protein backbones with the workflow described in Mueller et al. (2023). The implementation of the maab-pipeline (Johnson et al., 2017a) in the R-package “ragp” (Dragićević et al., 2019) was used to classify the putative AGP backbones into 24 distinct MAAB classes (Johnson et al., 2017a, Dataset S5) according to their motif counts and amino acid bias towards PAST. AG-peptides were excluded from the analysis by a length filter of ≥ 90 amino acids. Sequences containing at least four (STAGV)P dipeptides at a maximal distance of four amino acids plus an annotated pfam domain were investigated as chimeric AGPs (Dataset S6). The identification workflow from Mueller et al. (2023) was used for CAZyme analysis. Filtered GT and GH candidates were then aligned using MAFFT (Katoh et al., 2019; Katoh and Standley, 2013) with subsequent tree inference using the IQ-Tree webserver (Trifinopoulos et al., 2016). A maximum-likelihood approach with 1000 ultrafast bootstrap replicates (Minh et al., 2013) was used to generate the phylogenetic trees. The built-in model finder (Kalyaanamoorthy et al., 2017) was used to determine the optimal evolutionary model used for tree inference (for details see Datasets S4). *Potamogeton acutifolia* (Potamogetonaceae), a submersed alismatid monocot and sister group to the Zosteraceae (Chen et al., 2022) was used for reference.

### Analysis of differentially expressed genes

We investigated up- and down-regulated genes of importance in AGP biosynthesis under saline conditions using published raw datasets for *Thalassia hemprichii* (Shen et al., 2022; control: SRS13379160, SRS13379149, SRS13379148; salinity 48h / 45 PSU: SRS13379187, SRS13379189,

SRS13379188) and *Cymodocea nodosa* (Malandrakis et al., 2017; control: SRS847045, SRS847048, SRS847053; salinity 24h / 50 PSU: SRS846996, SRS847049, SRS847051). Datasets were analyzed on the Galaxy web platform (usegalaxy.org; Afgan et al., 2018) using the workflow of Bérénice Batut et al. (2024). The FASTQ reads were trimmed with Cutadapt (Martin, 2011) for removal of adapter sequences and reads with high uncertainty (default settings plus minimum read length: 20; quality cutoff: 20). The trimmed reads were mapped on the genomes of *Cymodocea nodosa* (Ma et al., 2024) and *Thalassia testudinum* (Ma et al., 2024) using STAR (Dobin et al., 2013). Differentially expressed genes were identified using DESeq2 (Love et al., 2014). Adjusted p-values below 0.05 were treated as significant hits. Fold change (FC) of > 2 and < 0.5 were interpreted as significantly up- or downregulated, respectively. Generated datasets are available as Dataset S7-S9.

## Acknowledgments

Thank you to the Malgana Aboriginal Corporation for permission to conduct seagrass research on Gathaagudu (Shark Bay). We thank Prof. Dr. Joseph A. Borg (University of Malta) for kind collection of *Cymodocea nodosa* plants used for biochemical investigations.

## Author contributions

LP and BC planned and designed the research. LP, EAS and CMD collected the plant material. LP performed the extractions, biochemical experiments and bioinformatics. TBHR provided access to genomic information. All authors analyzed and discussed the data. LP and BC drafted the manuscript. All authors discussed and finalized the paper. All authors read and approved the final manuscript.

## Supplemental data

The following materials are available in the online version of this article.

**Supplemental Table S1.** Yields of the extracted fractions

**Supplemental Table S2.** Monosaccharide composition (% mol/mol) for AGPs of the investigated seagrass species

**Supplemental Table S3.** Detailed list of seagrass origin and collection information

**Supplemental Figure S1.** Gas-chromatograms of the linkage-type analysis for *P. sinuosa* carboxy-reduced AGP (A) and oxalic-acid treated sample (B). The peak around 38.0 min represents the internal standard *myo*-inositol.

**Supplemental Figure S2.** Linkage-types of the different carboxy-reduced arabinogalactan-protein samples as inferred by methylation analysis. The relative amounts of galactoses in four distinct linkage-types (A: t-Galp, B:1,3-Galp, C: 1,6 Galp and D: 1,3,6-Galp) is shown prior and after acid degradation.

**Supplemental Figure S3.** BUSCO scores for assessment of genome and transcriptome quality. The reference database eukaryota_odb10 was used with BUSCO version 5.7.1.

**Dataset S1.** FASTA formatted files containing the sequences used for generation of the phylogenetic trees

**Dataset S2.** Alignment files used for phylogenetic analyses

**Dataset S3.** Tree files used for phylogenetic analyses

**Dataset S4.** Annotated trees for phylogenetic analyses

**Dataset S5.** FASTA formatted files containing the annotated AGP sequences from the maab analyses

**Dataset S6.** FASTA formatted files containing the annotated chimeric AGP sequences

**Dataset S7.** Differentially expressed genes (DEGs) of *Cymodocea nodosa* (Cymno) and *Thalassia hemprichii* (Thate, mapped on the genome of *Thalassia testudinum*) as summary tables

**Dataset S8.** Results of the DBCAN2 analysis of the DEGs of *Cymodocea nodosa* (Cymno) and *Thalassia hemprichii* (Thate, mapped on the genome of *Thalassia testudinum*) as summary tables

**Dataset S9.** Significantly up- or downregulated protein sequences of *Cymodocea nodosa* (Cymno) and *Thalassia hemprichii* (Thate, mapped on the genome of *Thalassia testudinum*) in FASTA format.

## Data availability

All relevant data can be found within the manuscript and its supporting materials.

## References

Ackerman JD. Sexual Reproduction of Seagrasses: Pollination in the Marine Context. In AWD Larkum, RJ Orth, CM Duarte, eds, Seagrasses: Biology, Ecology and Conservation. Springer Netherlands, Dordrecht, 2006, pp. 89–109.

Afgan E, Baker D, Batut B, van den Beek M, Bouvier D, Cech M, Chilton J, Clements D, Coraor N, Grüning BA, Guerler A, Hillman-Jackson J, Hiltemann S, Jalili V, Rasche H, Soranzo N, Goecks J, Taylor J, Nekrutenko A, Blankenberg D. The Galaxy platform for accessible, reproducible and collaborative biomedical analyses: 2018 update. Nucleic Acids Research. 2018:46: W537–W544. 10.1093/nar/gky379

Aires T, Marbà N, Cunha RL, Kendrick GA, Di Walker, Serrão EA, Duarte CM, Arnaud-Haond S. Evolutionary history of the seagrass genus Posidonia. Marine Ecology Progress Series. 2011:421: 117–130. 10.3354/meps08879

Aquino RS, Grativol C, Mourão PAS. Rising from the sea: correlations between sulfated polysaccharides and salinity in plants. PloS ONE. 2011:6: e18862. 10.1371/journal.pone.0018862

Aquino RS, Landeira-Fernandez AM, Valente AP, Andrade LR, Mourão PAS. Occurrence of sulfated galactans in marine angiosperms: evolutionary implications. Glycobiology. 2005:15: 11–20. 10.1093/glycob/cwh138

Bartels D, Baumann A, Maeder M, Geske T, Heise EM, Schwartzenberg K von, Classen B. Evolution of plant cell wall: Arabinogalactan-proteins from three moss genera show structural differences compared to seed plants. Carbohydrate Polymers. 2017:163: 227–235. 10.1016/j.carbpol.2017.01.043

Bartels D, Classen B. Structural investigations on arabinogalactan-proteins from a lycophyte and different monilophytes (ferns) in the evolutionary context. Carbohydrate Polymers. 2017:172: 342–351. 10.1016/j.carbpol.2017.05.031

Basu D, Tian L, Debrosse T, Poirier E, Emch K, Herock H, Travers A, Showalter AM. Glycosylation of a Fasciclin-Like Arabinogalactan-Protein (SOS5) Mediates Root Growth and Seed Mucilage Adherence via a Cell Wall Receptor-Like Kinase (FEI1/FEI2) Pathway in Arabidopsis. PloS ONE. 2016:11: e0145092. 10.1371/journal.pone.0145092

Basu D, Tian L, Wang W, Bobbs S, Herock H, Travers A, Showalter AM. A small multigene hydroxyproline-O-galactosyltransferase family functions in arabinogalactan-protein glycosylation, growth and development in Arabidopsis. BMC Plant Biology. 2015a:15: 295. 10.1186/s12870-015-0670-7

Basu D, Wang W, Ma S, DeBrosse T, Poirier E, Emch K, Soukup E, Tian L, Showalter AM. Two Hydroxyproline Galactosyltransferases, GALT5 and GALT2, Function in Arabinogalactan-Protein Glycosylation, Growth and Development in Arabidopsis. PloS ONE. 2015b:10: e0125624. 10.1371/journal.pone.0125624

Baumann A, Pfeifer L, Classen B. Arabinogalactan-proteins from non-coniferous gymnosperms have unusual structural features. Carbohydrate Polymers. 2021:261: 117831. 10.1016/j.carbpol.2021.117831

Bayer PE, Fraser MW, Martin BC, Petereit J, Severn-Ellis AA, Sinclair EA, Batley J, Kendrick GA, Edwards D. Not all pathways are the same – unique adaptations to submerged environments emerge from comparative seagrass genomics. 10.1101/2022.11.22.517588

Bérénice Batut, Mallory Freeberg, Mo Heydarian, Anika Erxleben, Pavankumar Videm, Clemens Blank, Maria Doyle, Nicola Soranzo, Peter van Heusden, Lucille Delisle. (2024) Reference-based RNA-Seq data analysis (Galaxy Training Materials), https://training.galaxyproject.org/training-material/topics/transcriptomics/tutorials/ref-based/tutorial.html.

Blakeney AB, Harris PJ, Henry RJ, Stone BA. A simple and rapid preparation of alditol acetates for monosaccharide analysis. Carbohydrate Research. 1983:113: 291–299. 10.1016/0008-6215(83)88244-5

Booth MW, Breed MF, Kendrick GA, Bayer PE, Severn-Ellis AA, Sinclair EA. Tissue-specific transcriptome profiles identify functional differences key to understanding whole plant response to life in variable salinity. Biology Open. 2023:11. 10.1242/bio.059147

Borg M, Krueger-Hadfield SA, Destombe C, Collén J, Lipinska A, Coelho SM. Red macroalgae in the genomic era. New Phytologist. 2023:240: 471–488. 10.1111/nph.19211

Chen L-Y, Lu B, Morales-Briones DF, Moody ML, Liu F, Hu G-W, Huang C-H, Chen J-M, Wang Q-F. Phylogenomic Analyses of Alismatales Shed Light into Adaptations to Aquatic Environments. Molecular Biology and Evolution. 2022:39:msac079, 10.1093/molbev/msac079

Ciancia M, Matulewicz MC, Tuvikene R. Structural Diversity in Galactans From Red Seaweeds and Its Influence on Rheological Properties. Frontiers in Plant Science. 2020:11: 559986. 10.3389/fpls.2020.559986

Coimbra S, Almeida J, Junqueira V, Costa ML, Pereira LG. Arabinogalactan proteins as molecular markers in Arabidopsis thaliana sexual reproduction. Journal of Experimental Botany. 2007:58: 4027–4035. 10.1093/jxb/erm259

Costa M, Pereira AM, Pinto SC, Silva J, Pereira LG, Coimbra S. In silico and expression analyses of fasciclin-like arabinogalactan proteins reveal functional conservation during embryo and seed development. Plant Reproduction. 2019:32: 353–370. 10.1007/s00497-019-00376-7

Cramer GR, Läuchli A, Polito VS. Displacement of Ca^2+^ by Na^+^ from the plasmalemma of root cells: a primary response to salt stress? Plant Physiology. 1985:79: 207–211. 10.1104/pp.79.1.207

Da Silva SL, Carvalho R de, Magalhães KM. Chromosomal evolution in seagrasses: Is the chromosome number decreasing? Aquatic Botany. 2021:173: 103410. 10.1016/j.aquabot.2021.103410

Davies P, Morvan C, Sire O, Baley C. Structure and properties of fibres from sea-grass (Zostera marina). Journal of Materials Science. 2007:42: 4850–4857. 10.1007/s10853-006-0546-1

Denoeud F, Godfroy O, Cruaud C, Heesch S, Nehr Z, Tadrent N, Couloux A, Brillet-Guéguen L, Delage L, Mckeown D, et al. Evolutionary genomics of the emergence of brown algae as key components of coastal ecosystems. Cell. 2024:187: 6943–6965.e39. 10.1016/j.cell.2024.10.049

Dilokpimol A, Geshi N. Arabidopsis thaliana glucuronosyltransferase in family GT14. Plant Signaling & Behavior. 2014:9: e28891. 10.4161/psb.28891

Dobin A, Davis CA, Schlesinger F, Drenkow J, Zaleski C, Jha S, Batut P, Chaisson M, Gingeras TR. STAR: ultrafast universal RNA-seq aligner. Bioinformatics (Oxford, England). 2013:29: 15–21. 10.1093/bioinformatics/bts635

Dragićević MB, Paunović DM, Bogdanović MD, Todorović SI, Simonović AD. ragp: Pipeline for mining of plant hydroxyproline-rich glycoproteins with implementation in R. Glycobiology.2019:11: cwz072. 10.1093/glycob/cwz072

Duarte CM, Losada IJ, Hendriks IE, Mazarrasa I, Marbà N. The role of coastal plant communities for climate change mitigation and adaptation. Nature Climate Change. 2013:3: 961– 968. 10.1038/nclimate1970

Duarte CM, Middelburg JJ, Caraco N. Major role of marine vegetation on the oceanic carbon cycle. Biogeosciences. 2005:2: 1–8. 10.5194/bg-2-1-2005

Eder M, Tenhaken R, Driouich A, Lütz-Meindl U. Occurrence and characterization of arabinogalactan-like proteins and hemicelluloses in *Micrasterias* (Streptophyta). Journal of Phycology. 2008:44: 1221–1234. 10.1111/j.1529-8817.2008.00576.x

Ellis M, Egelund J, Schultz CJ, Bacic A. Arabinogalactan-proteins: key regulators at the cell surface? Plant Physiology. 2010:153: 403–419.

Eudes A, Mouille G, Thévenin J, Goyallon A, Minic Z, Jouanin L. Purification, cloning and functional characterization of an endogenous beta-glucuronidase in *Arabidopsis thaliana*. Plant & Cell Physiology. 2008:49: 1331–1341. 10.1093/pcp/pcn108

Fincher GB, Stone BA, Clarke AE. Arabinogalactan-proteins: structure, biosynthesis, and function. Annual Review of Plant Biology, Palo Alto. 1983:34: 47–70. 10.1146/annurev.pp.34.060183.000403

Franková L, Fry SC. Biochemistry and physiological roles of enzymes that ‘cut and paste’ plant cell-wall polysaccharides. Journal of Experimental Botany. 2013:64: 3519–3550. 10.1093/jxb/ert201

Fu H, Yadav MP, Nothnagel EA. *Physcomitrella patens* arabinogalactan proteins contain abundant terminal 3-*O*-methyl-L-rhamnosyl residues not found in angiosperms. Planta. 2007:226: 1511– 1524. 10.1007/s00425-007-0587-y

Galili T. dendextend: an R package for visualizing, adjusting and comparing trees of hierarchical clustering. Bioinformatics (Oxford, England). 2015:31: 3718–3720. 10.1093/bioinformatics/btv428

Gille S, Sharma V, Baidoo EEK, Keasling JD, Scheller HV, Pauly M. Arabinosylation of a Yariv-precipitable cell wall polymer impacts plant growth as exemplified by the Arabidopsis glycosyltransferase mutant ray1. Molecular Plant: 2013:6: 1369–1372. 10.1093/mp/sst029

Gloaguen V, Brudieux V, Closs B, Barbat A, Krausz P, Sainte-Catherine O, Kraemer M, Maes E, Guerardel Y. Structural characterization and cytotoxic properties of an apiose-rich pectic polysaccharide obtained from the cell wall of the marine phanerogam *Zostera marina*. Journal of Natural Products. 2010:73: 1087–1092. 10.1021/np100092c

Grant GT, Morris ER, Rees DA, Smith PJ, Thom D. Biological interactions between polysaccharides and divalent cations: The egg-box model. FEBS Letters. 1973:32: 195–198. 10.1016/0014-5793(73)80770-7

Harris PJ, Henry RJ, Blakeney AB, Stone BA. An improved procedure for the methylation analysis of oligosaccharides and polysaccharides. Carbohydrate Research. 1984:127: 59–73. 10.1016/0008-6215(84)85106-X

He J, Zhao H, Cheng Z, Ke Y, Liu J, Ma H. Evolution Analysis of the Fasciclin-Like Arabinogalactan Proteins in Plants Shows Variable Fasciclin-AGP Domain Constitutions. International Journal of Molecular Sciences. 2019:20. 10.3390/ijms20081945

Hervé C, Siméon A, Jam M, Cassin A, Johnson KL, Salmeán AA, Willats WGT, Doblin MS, Bacic A, Kloareg B. Arabinogalactan proteins have deep roots in eukaryotes: identification of genes and epitopes in brown algae and their role in Fucus serratus embryo development. New Phytologist. 2016:209: 1428–1441. 10.1111/nph.13786

Hoegh-Guldberg O, Cai R, Poloczanska ES, Brewer PG, Sundby S, Hilmi K, Fabry VJ, Jung S, Skirving W, Stone DA, et al. The Ocean. In: Climate Change 2014: Impacts, Adaptation, and Vulnerability. Part B: Regional Aspects. Contribution of Working Group II to the Fifth Assessment Report of the Intergovernmental Panel on Climate Change. Barros, V.R., C.B. Field, D.J. Dokken, M.D. Mastrandrea, K.J. Mach, T.E. Bilir, M. Chatterjee, K.L. Ebi, Y.O. Estrada, R.C. Genova, B. Girma, E.S. Kissel, A.N. Levy, S. MacCracken, P.R. Mastrandrea, and L.L. White, eds., Cambridge University Press, Cambridge, United Kingdom and New York, NY, USA, pp. 1655–1731.

Imaizumi C, Tomatsu H, Kitazawa K, Yoshimi Y, Shibano S, Kikuchi K, Yamaguchi M, Kaneko S, Tsumuraya Y, Kotake T. Heterologous expression and characterization of an Arabidopsis β-l-arabinopyranosidase and α-d-galactosidases acting on β-l-arabinopyranosyl residues. Journal of Experimental Botany. 2017:68: 4651–4661. 10.1093/jxb/erx279

Ismael M, Charras Q, Leschevin M, Herfurth D, Roulard R, Quéro A, Rusterucci C, Domon J-M, Jungas C, Vermerris W, Rayon C. Seasonal Variation in Cell Wall Composition and Carbohydrate Metabolism in the Seagrass Posidonia oceanica Growing at Different Depths. *Plants* (Basel, Switzerland). 2023:12: 3155. 10.3390/plants12173155

Ito K, Fukuoka K, Nishigaki N, Hara K, Yoshimi Y, Kuki H, Takahashi D, Tsumuraya Y, Kotake T. Structural features conserved in subclass of type II arabinogalactan. *Plant Biotechnology* (Tokyo, Japan). 2020:37: 459–463. 10.5511/plantbiotechnology.20.0721a

Iwai H, Masaoka N, Ishii T, Satoh S. A pectin glucuronyltransferase gene is essential for intercellular attachment in the plant meristem. Proceedings of the National Academy of Sciences of the United States of America. 2002:99: 16319–16324. 10.1073/pnas.252530499

Johnson KL, Cassin AM, Lonsdale A, Bacic A, Doblin MS, Schultz CJ. Pipeline to Identify Hydroxyproline-Rich Glycoproteins. Plant Physiology. 2017a:174: 886–903. 10.1104/pp.17.00294

Johnson KL, Cassin AM, Lonsdale A, Wong GK-S, Soltis DE, Miles NW, Melkonian M, Melkonian B, Deyholos MK, Leebens-Mack J, et al. Insights into the Evolution of Hydroxyproline-Rich Glycoproteins from 1000 Plant Transcriptomes. Plant Physiology. 2017b:174: 904–921. 10.1104/pp.17.00295

Johnson KL, Jones BJ, Bacic A, Schultz CJ. The fasciclin-like arabinogalactan proteins of Arabidopsis. A multigene family of putative cell adhesion molecules. Plant Physiology. 2003:133: 1911–1925. 10.1104/pp.103.031237

Kaal J, Serrano O, Del Río JC, Rencoret J. Radically different lignin composition in Posidonia species may link to differences in organic carbon sequestration capacity. Organic Geochemistry. 2018:124: 247–256. 10.1016/j.orggeochem.2018.07.017

Kalyaanamoorthy S, Minh BQ, Wong TKF, Haeseler A von, Jermiin LS. ModelFinder: fast model selection for accurate phylogenetic estimates. Nature Methods. 2017:14: 587–589. 10.1038/nmeth.4285

Katoh K, Rozewicki J, Yamada KD. MAFFT online service: multiple sequence alignment, interactive sequence choice and visualization. Briefings in Bioinformatics. 2019:20: 1160–1166. 10.1093/bib/bbx108

Katoh K, Standley DM. MAFFT multiple sequence alignment software version 7: improvements in performance and usability. Molecular Biology and Evolution. 2013:30: 772–780. 10.1093/molbev/mst010

Kikuchi A, Hara K, Yoshimi Y, Soga K, Takahashi D, Kotake T. *In vivo* structural modification of type II arabinogalactans with fungal endo-β-1, 6-galactanase in Arabidopsis. Frontiers in Plant Science. 2022:13: 1010492. 10.3389/fpls.2022.1010492

Kim JB, Carpita NC. Changes in Esterification of the Uronic Acid Groups of Cell Wall Polysaccharides during Elongation of Maize Coleoptiles. Plant Physiology. 1992:98: 646–653. 10.1104/pp.98.2.646

Knoch E, Dilokpimol A, Tryfona T, Poulsen CP, Xiong G, Harholt J, Petersen BL, Ulvskov P, Hadi MZ, Kotake T, Tsumuraya Y, Pauly M, Dupree P, Geshi N. A β-glucuronosyltransferase from Arabidopsis thaliana involved in biosynthesis of type II arabinogalactan has a role in cell elongation during seedling growth. The Plant Journal. 2013:76: 1016–1029. 10.1111/tpj.12353

Knox JP, Linstead PJ, Cooper JPC, Roberts K. Developmentally regulated epitopes of cell surface arabinogalactan proteins and their relation to root tissue pattern formation. The Plant Journal. 1991:1: 317–326. 10.1046/j.1365-313x.1991.t01-9-00999.x

Kolsi RBA, Fakhfakh J, Krichen F, Jribi I, Chiarore A, Patti FP, Blecker C, Allouche N, Belghith H, Belghith K. Structural characterization and functional properties of antihypertensive *Cymodocea nodosa* sulfated polysaccharide. Carbohydrate Polymers. 2016:151: 511–522. 10.1016/j.carbpol.2016.05.098

Kotake T, Dina S, Konishi T, Kaneko S, Igarashi K, Samejima M, Watanabe Y, Kimura K, Tsumuraya Y. Molecular cloning of a {beta}-galactosidase from radish that specifically hydrolyzes {beta}-(1-3)- and {beta}-(1-6)-galactosyl residues of Arabinogalactan protein. Plant Physiology. 2005:138: 1563–1576. 10.1104/pp.105.062562

Kotake T, Tsuchiya K, Aohara T, Konishi T, Kaneko S, Igarashi K, Samejima M, Tsumuraya Y. An alpha-L-arabinofuranosidase/beta-D-xylosidase from immature seeds of radish (Raphanus sativus L.). Journal of Experimental Botany. 2006:57: 2353–2362. 10.1093/jxb/erj206

Lahaye PA, Epstein E. Salt toleration by plants: enhancement with calcium. *Science* (New York, N.Y.). 1969:166: 395–396. 10.1126/science.166.3903.395

Lamport DTA. Climate change and the single cell. Academia Biology. 2024:2: 4. 10.20935/AcadBiol7421

Lamport DTA, Tan L, Held M, Kieliszewski MJ. Phyllotaxis Turns Over a New Leaf-A New Hypothesis. International Journal of Molecular Sciences. 2020:21: 1145. 10.3390/ijms21031145

Lamport DTA, Varnai P, Seal CE. Back to the future with the AGP-Ca^2+^ flux capacitor. Annals of Botany. 2014:114: 1069–1085. 10.1093/aob/mcu161

Lamport DTA, Várnai P. Periplasmic arabinogalactan glycoproteins act as a calcium capacitor that regulates plant growth and development. New Phytologist. 2013:197: 58–64. 10.1111/nph.12005

Larkum AWD, Waycott M, Conran JG. Evolution and Biogeography of Seagrasses. In AWD Larkum, GA Kendrick, PJ Ralph, eds, Seagrasses of Australia. 2018. Springer International Publishing, Cham, pp. 3–29.

Lee H, Golicz AA, Bayer PE, Jiao Y, Tang H, Paterson AH, Sablok G, Krishnaraj RR, Chan C-KK, Batley J, et al. The Genome of a Southern Hemisphere Seagrass Species (*Zostera muelleri*). Plant Physiology. 2016:172: 272–283. 10.1104/pp.16.00868

Lee H, Golicz AA, Bayer PE, Severn-Ellis AA, Chan C-KK, Batley J, Kendrick GA, Edwards D. Genomic comparison of two independent seagrass lineages reveals habitat-driven convergent evolution. Journal of Experimental Botany. 2018:69: 3689–3702. 10.1093/jxb/ery147

Les DH, Cleland MA, Waycott M. Phylogenetic Studies in Alismatidae, II: Evolution of Marine Angiosperms (Seagrasses) and Hydrophily. Systematic Botany. 1997:22: 443. 10.2307/2419820

Leszczuk A, Kalaitzis P, Kulik J, Zdunek A. Review: structure and modifications of arabinogalactan proteins (AGPs). BMC Plant Biology. 2023:23: 45. 10.1186/s12870-023-04066-5

Li P, Li Y-J, Zhang F-J, Zhang G-Z, Jiang X-Y, Yu H-M, Hou B-K. The *Arabidopsis* UDP-glycosyltransferases UGT79B2 and UGT79B3, contribute to cold, salt and drought stress tolerance via modulating anthocyanin accumulation. The Plant Journal. 2017:89: 85–103. 10.1111/tpj.13324

Lin Y, Lian L, Zhu Y, Wang L, Li H, Zheng Y, Cai Q, He W, Xie H, Wei Y, et al. Characterization and expression analysis of the glycosyltransferase 64 family in rice (Oryza sativa). Gene. 2022:838: 146708. 10.1016/j.gene.2022.146708

Lischer HEL, Shimizu KK. Reference-guided de novo assembly approach improves genome reconstruction for related species. BMC Bioinformatics. 2017:18: 474. 10.1186/s12859-017-1911-6

Lopez-Hernandez F, Tryfona T, Rizza A, Yu XL, Harris MOB, Webb AAR, Kotake T, Dupree P. Calcium Binding by Arabinogalactan Polysaccharides Is Important for Normal Plant Development. The Plant Cell. 2020:32: 3346–3369. 10.1105/tpc.20.00027

Love MI, Huber W, Anders S. Moderated estimation of fold change and dispersion for RNA-seq data with DESeq2. Genome Biology. 2014:15: 550. 10.1186/s13059-014-0550-8

Lv Y, Shan X, Zhao X, Cai C, Zhao X, Lang Y, Zhu H, Yu G. Extraction, Isolation, Structural Characterization and Anti-Tumor Properties of an Apigalacturonan-Rich Polysaccharide from the Sea Grass *Zostera caespitosa* Miki. Marine Drugs. 2015:13: 3710–3731. 10.3390/md13063710

Ma X, Vanneste S, Chang J, Ambrosino L, Barry K, Bayer T, Bobrov AA, Boston L, Campbell JE, Chen H, et al. Seagrass genomes reveal ancient polyploidy and adaptations to the marine environment. Nature Plants. 2024:10: 240–255. 10.1038/s41477-023-01608-5

Ma Y, Johnson K. Arabinogalactan proteins - Multifunctional glycoproteins of the plant cell wall. The Cell Surface. 2023:9: 100102. 10.1016/j.tcsw.2023.100102

Ma Y, Yan C, Li H, Wu W, Liu Y, Wang Y, Chen Q, Ma H. Bioinformatics Prediction and Evolution Analysis of Arabinogalactan Proteins in the Plant Kingdom. Frontiers in Plant Science. 2017:8: 66. 10.3389/fpls.2017.00066

MacMillan CP, Mansfield SD, Stachurski ZH, Evans R, Southerton SG. Fasciclin-like arabinogalactan proteins: specialization for stem biomechanics and cell wall architecture in Arabidopsis and Eucalyptus. The Plant Journal. 2010:62: 689–703. 10.1111/j.1365-313x.2010.04181.x

Malandrakis E, Dadali O, Kavouras M, Danis T, Panagiotaki P, Miliou H, Tsioli S, Orfanidis S, Küpper FC, Exadactylos A. Identification of the abiotic stress-related transcription in little Neptune grass *Cymodocea nodosa* with RNA-seq. Marine Genomics. 2017:34: 47–56. 10.1016/j.margen.2017.03.005

Martin M. Cutadapt removes adapter sequences from high-throughput sequencing reads. EMBnet.journal. 2011:17: 10. 10.14806/ej.17.1.200

Mazéas L, Bouguerba-Collin A, Cock JM, Denoeud F, Godfroy O, Brillet-Guéguen L, Barbeyron T, Lipinska AP, Delage L, Corre E, et al. Candidate genes involved in biosynthesis and degradation of the main extracellular matrix polysaccharides of brown algae and their probable evolutionary history. BMC Genomics. 2024:25: 950. 10.1186/s12864-024-10811-3

Minh BQ, Nguyen MAT, Haeseler A. Ultrafast approximation for phylogenetic bootstrap. Molecular Biology and Evolution. 2013:30: 1188–1195. 10.1093/molbev/mst024

Mizukami AG, Inatsugi R, Jiao J, Kotake T, Kuwata K, Ootani K, Okuda S, Sankaranarayanan S, Sato Y, Maruyama D, et al. The AMOR Arabinogalactan Sugar Chain Induces Pollen-Tube Competency to Respond to Ovular Guidance. Current Biology. 2016:26: 1091–1097. 10.1016/j.cub.2016.02.040

Mueller K-K, Pfeifer L, Schuldt L, Szövényi P, Vries S de, Vries J de, Johnson KL, Classen B. Fern cell walls and the evolution of arabinogalactan proteins in streptophytes. The Plant Journal. 2023:114: 875–894. 10.1111/tpj.16178

Nibbering P, Castilleux R, Wingsle G, Niittylä T. CAGEs are Golgi-localized GT31 enzymes involved in cellulose biosynthesis in Arabidopsis. The Plant Journal. 2022:110: 1271–1285. 10.1111/tpj.15734

Nibbering P, Petersen BL, Motawia MS, Jørgensen B, Ulvskov P, Niittylä T. Golgi-localized exo-β1,3-galactosidases involved in cell expansion and root growth in Arabidopsis. The Journal of Biological Chemistry. 2020:295: 10581–10592. 10.1074/jbc.ra120.013878

Nothnagel EA. Proteoglycans and related components in plant cells. International Review of Cytology. 1997:174: 195–291. 10.1016/s0074-7696(08)62118-x

Olmos E, La García De Garma J, Gomez-Jimenez MC, Fernandez-Garcia N. Arabinogalactan Proteins Are Involved in Salt-Adaptation and Vesicle Trafficking in Tobacco by-2 Cell Cultures. Frontiers in Plant Science. 2017:8: 1092. 10.3389/fpls.2017.01092

Olsen JL, Rouzé P, Verhelst B, Lin Y-C, Bayer T, Collen J, Dattolo E, Paoli E de, Dittami S, Maumus F, et al. The genome of the seagrass Zostera marina reveals angiosperm adaptation to the sea. Nature. 2016:530: 331–335. 10.1038/nature16548

Papenbrock J. Highlights in Seagrasses’ Phylogeny, Physiology, and Metabolism: What Makes Them Special? 2012:103892. 10.5402/2012/103892

Pelloux J, Rustérucci C, Mellerowicz EJ. New insights into pectin methylesterase structure and function. Trends in Plant Science. 2007:12: 267–277. 10.1016/j.tplants.2007.04.001

Pereira AM, Lopes AL, Coimbra S. Arabinogalactan Proteins as Interactors along the Crosstalk between the Pollen Tube and the Female Tissues. Frontiers in Plant Science. 2016:7: 1895. 10.3389/fpls.2016.01895

Pereira AM, Pereira LG, Coimbra S. Arabinogalactan proteins: rising attention from plant biologists. Plant Reproduction. 2015:28: 1–15. 10.1007/s00497-015-0254-6

Pervaiz T, Liu T, Fang X, Ren Y, Li X, Liu Z, Fiaz M, Fang J, Shangguan L. Identification of GH17 gene family in Vitis vinifera and expression analysis of GH17 under various adversities. Physiology and Molecular Biology of plants. 2021:27: 1423–1436. 10.1007/s12298-021-01014-1

Pfeifer L. Enigmatic Apiogalacturonans – What Do We Know About This Group of Pectic Polysaccharides? In JA Roberts, ed, Annual Plant Reviews online. Wiley, 2023, pp. 1–30. 10.1002/9781119312994.apr0803

Pfeifer L, Classen B. The Cell Wall of Seagrasses: Fascinating, Peculiar and a Blank Canvas for Future Research. Frontiers in Plant Science. 2020a:11: 588754. 10.3389/fpls.2020.588754

Pfeifer L, Classen B. Validation of a Rapid GC-MS Procedure for Quantitative Distinction between 3-*O*-Methyl- and 4-*O*-Methyl-Hexoses and Its Application to a Complex Carbohydrate Sample. Separations. 2020b:7: 42. 10.3390/separations7030042

Pfeifer L, Mueller K-K, Utermöhlen J, Erdt F, Zehge JBJ, Schubert H, Classen B. The cell walls of different *Chara* species are characterized by branched galactans rich in 3-*O*-methylgalactose and absence of AGPs. Physiologia Plantarum. 2023:175: e13989. 10.1111/ppl.13989

Pfeifer L, Shafee T, Johnson KL, Bacic A, Classen B. Arabinogalactan-proteins of *Zostera marina* L. contain unique glycan structures and provide insight into adaption processes to saline environments. Scientific Reports. 2020:10: 8232. 10.1038/s41598-020-65135-5

Pfeifer L, Utermöhlen J, Happ K, Permann C, Holzinger A, Schwartzenberg K von, Classen B. Search for evolutionary roots of land plant arabinogalactan-proteins in charophytes: presence of a rhamnogalactan-protein in *Spirogyra pratensis* (Zygnematophyceae). The Plant Journal. 2022a:109: 568–584. 10.1111/tpj.15577

Pfeifer L, van Erven G, Sinclair EA, Duarte CM, Kabel MA, Classen B. Profiling the cell walls of seagrasses from A (*Amphibolis*) to Z (*Zostera*). BMC Plant Biology. 2022b:22: 63. 10.1186/s12870-022-03447-6

Přerovská T, Henke S, Bleha R, Spiwok V, Gillarová S, Yvin J-C, Ferrières V, Nguema-Ona E, Lipovová P. Arabinogalactan-like Glycoproteins from Ulva lactuca (Chlorophyta) Show Unique Features Compared to Land Plants AGPs. Journal of Phycology. 2021:57: 619–635. 10.1111/jpy.13121

Rennie EA, Hansen SF, Baidoo EEK, Hadi MZ, Keasling JD, Scheller HV. Three members of the Arabidopsis glycosyltransferase family 8 are xylan glucuronosyltransferases. Plant Physiology. 2012:159: 1408–1417. 10.1104/pp.112.200964

Röthig T, Trevathan-Tackett SM, Voolstra CR, Ross C, Chaffron S, Durack PJ, Warmuth LM, Sweet M. Human-induced salinity changes impact marine organisms and ecosystems. Global Change Biology. 2023:29: 4731–4749. 10.1111/gcb.16859

Ruprecht C, Bartetzko MP, Senf D, Dallabernadina P, Boos I, Andersen MCF, Kotake T, Knox JP, Hahn MG, Clausen MH, Pfrengle F. A Synthetic Glycan Microarray Enables Epitope Mapping of Plant Cell Wall Glycan-Directed Antibodies. Plant Physiology. 2017:175: 1094–1104. 10.1104/pp.17.00737

Sandoval-Gil JM, Ruiz JM, Marín-Guirao L. Advances in understanding multilevel responses of seagrasses to hypersalinity. Marine Environmental Research. 2023:183: 105809. 10.1016/j.marenvres.2022.105809

Sasaki Y, Uchimura Y, Kitahara K, Fujita K. Characterization of a GH36 α-D-Galactosidase Associated with Assimilation of Gum Arabic in *Bifidobacterium longum* subsp. *longum* JCM7052. Journal of Applied Glycoscience. 2021:68: 47–52. 10.5458/jag.jag.JAG-2021_0004

Seifert GJ. On the Potential Function of Type II Arabinogalactan O-Glycosylation in Regulating the Fate of Plant Secretory Proteins. Frontiers in Plant Science. 2020:11: 563735. 10.3389/fpls.2020.563735

Seifert GJ, Roberts K. The biology of arabinogalactan proteins. Annual Review of Plant Biology. 2007:58: 137–161. 10.1146/annurev.arplant.58.032806.103801

Shafee T, Bacic A, Johnson K. Evolution of Sequence-Diverse Disordered Regions in a Protein Family: Order within the Chaos. Molecular Biology and Evolution. 2020:37: 2155–2172. 10.1093/molbev/msaa096

Shen J, Wu Z, Yin L, Chen S, Cai Z, Geng X, Wang D. Physiological basis and differentially expressed genes in the salt tolerance mechanism of *Thalassia hemprichii*. Frontiers in Plant Science. 2022:13: 975251. 10.3389/fpls.2022.975251

Short F, Carruthers T, Dennison W, Waycott M. Global seagrass distribution and diversity: A bioregional model. Journal of Experimental Marine Biology and Ecology. 2007:350: 3–20. 10.1016/j.jembe.2007.06.012

Showalter AM, Basu D. Extensin and Arabinogalactan-Protein Biosynthesis: Glycosyltransferases, Research Challenges, and Biosensors. Frontiers in Plant Science. 2016:7: 814. 10.3389/fpls.2016.00814

Silva J, Ferraz R, Dupree P, Showalter AM, Coimbra S. Three Decades of Advances in Arabinogalactan-Protein Biosynthesis. Frontiers in Plant Science. 2020:11: 610377. 10.3389/fpls.2020.610377

Smallwood M, Yates EA, Willats WGT, Martin H, Knox JP. Immunochemical comparison of membrane-associated and secreted arabinogalactan-proteins in rice and carrot. Planta. 1996:198: 452–459. 10.1007/BF00620063

Smith PJ, O’Neill MA, Backe J, York WS, Peña MJ, Urbanowicz BR. Analytical Techniques for Determining the Role of Domain of Unknown Function 579 Proteins in the Synthesis of *O*-Methylated Plant Polysaccharides. SLAS technology. 2020:25: 345–355. 10.1177/2472630320912692

Sprockett DD, Piontkivska H, Blackwood CB. Evolutionary analysis of glycosyl hydrolase family 28 (GH28) suggests lineage-specific expansions in necrotrophic fungal pathogens. Gene. 2011:479: 29–36. 10.1016/j.gene.2011.02.009

Stegemann H, Stalder K. Determination of hydroxyproline. Clinica Chimica Acta. 1976:18: 267–273. 10.1016/0009-8981(67)90167-2

Strasser R, Seifert G, Doblin MS, Johnson KL, Ruprecht C, Pfrengle F, Bacic A, Estevez JM. Cracking the “Sugar Code”: A Snapshot of N- and O-Glycosylation Pathways and Functions in Plants Cells. Frontiers in Plant Science. 2021:12: 640919. 10.3389/fpls.2021.640919

Syed NFN, Zakaria MH, Bujang JS. Fiber Characteristics and Papermaking of Seagrass Using Hand-beaten and Blended Pulp. BioResources. 2016:11. 10.15376/biores.11.2.5358-5380

Tan L, Showalter AM, Egelund J, Hernandez-Sanchez A, Doblin MS, Bacic A. Arabinogalactan-proteins and the research challenges for these enigmatic plant cell surface proteoglycans. Frontiers in Plant Science. 2012:3: 140. 10.3389/fpls.2012.00140

Taylor RL, Conrad HE. Stoichiometric depolymerization of polyuronides and glycosaminoglycuronans to monosaccharides following reduction of their carbodiimide-activated carboxyl groups. Biochemistry. 1972:11: 1383–1388. 10.1021/bi00758a009

Temple H, Mortimer JC, Tryfona T, Yu X, Lopez-Hernandez F, Sorieul M, Anders N, Dupree P. Two members of the DUF579 family are responsible for arabinogalactan methylation in Arabidopsis. Plant Direct. 2019:3: e00117. 10.1002/pld3.117

Touchette BW. Seagrass-salinity interactions: Physiological mechanisms used by submersed marine angiosperms for a life at sea. Journal of Experimental Marine Biology and Ecology. 2007:350: 194–215. 10.1016/j.jembe.2007.05.037

Trifinopoulos J, Nguyen L-T, Haeseler A von, Minh BQ. W-IQ-TREE: a fast online phylogenetic tool for maximum likelihood analysis. Nucleic Acids Research. 2016:44: W232–5. 10.1093/nar/gkw256

Tuya F, Martínez-Pérez J, Fueyo Á, Bosch NE. Strong phylogenetic signal and models of trait evolution evidence phylogenetic niche conservatism for seagrasses. Journal of Ecology. 2024:112: 446–460. 10.1111/1365-2745.14232

Unsworth RKF, Cullen-Unsworth LC, Jones BLH, Lilley RJ. The planetary role of seagrass conservation. *Science* (New York, N.Y.). 2022:377: 609–613. 10.1126/science.abq6923

Waycott M, Biffin E, Les DH. Systematics and Evolution of Australian Seagrasses in a Global Context. In AWD Larkum, GA Kendrick, PJ Ralph, eds, Seagrasses of Australia. 2018, Springer International Publishing, Cham, pp. 129–154.

Wei T, Simko V. R package ‘corrplot’: Visualization of a Correlation Matrix, 2024, https://github.com/taiyun/corrplot.

Willats WG, Marcus SE, Knox JP. Generation of monoclonal antibody specific to (1-->5)-alpha-L-arabinan. Carbohydrate Research. 1998:308: 149–152. 10.1016/s0008-6215(98)00070-6

Wissler L, Codoñer FM, Gu J, Reusch TBH, Olsen JL, Procaccini G, Bornberg-Bauer E. Back to the sea twice: identifying candidate plant genes for molecular evolution to marine life. BMC Evolutionary Biology. 2011:11: 8. 10.1186/1471-2148-11-8

Xu S, Wang J, Guo Z, He Z, Shi S. Genomic Convergence in the Adaptation to Extreme Environments. Plant Communications. 2020:1: 100117. 10.1016/j.xplc.2020.100117

Yariv J, Lis H, Katchalski E. Precipitation of arabic acid and some seed polysaccharides by glycosylphenylazo dyes. The Biochemical Journal. 1967:105: 1C–2C. 10.1042/bj1050001c

Yates EA, Valdor JF, Haslam SM, Morris HR, Dell A, Mackie W, Knox JP. Characterization of carbohydrate structural features recognized by anti-arabinogalactan-protein monoclonal antibodies. Glycobiology. 1996:6: 131–139. 10.1093/glycob/6.2.131

Zhong H, Läuchli A. Changes of Cell Wall Composition and Polymer Size in Primary Roots of Cotton Seedlings Under High Salinity. Journal of Experimental Botany. 1993:44: 773–778. 10.1093/jxb/44.4.773

Zhong R, Ye Z-H. Unraveling the functions of glycosyltransferase family 47 in plants. Trends in Plant Science. 2003:8: 565–568. 10.1016/j.tplants.2003.10.003

